# Neutrophil stalling does not mediate the increase in tau phosphorylation and the cognitive impairment associated with high salt diet

**DOI:** 10.1101/2025.02.27.640593

**Authors:** Sung-Ji Ahn, Benjamin Goya, Christian Bertomo, Rose Sciortino, Gianfranco Racchumi, Lidia Garcia Bonilla, Josef Anrather, Costantino Iadecola, Giuseppe Faraco

**Affiliations:** Feil Family Brain and Mind Research Institute, Weill Cornell Medicine, New York, NY 10065, USA

**Keywords:** High salt diet, capillary stalling, neutrophils, cognitive impairment, tau phosphorylation

## Abstract

High dietary salt intake has powerful effects on cerebral blood vessels and has emerged as a risk factor for stroke and cognitive impairment. In mice, high salt diet (HSD) leads to reduced cerebral blood flow (CBF), tau hyperphosphorylation and cognitive dysfunction. However, it is still unclear whether the reduced CBF is responsible for the effects of HSD on tau and cognition. Capillary stalling has emerged as a cause of CBF reduction and cognitive impairment in models of Alzheimer’s disease and diabetes. Therefore, we tested the hypothesis that capillary stalling also contributes to the CBF reduction and cognitive impairment in HSD. Using two-photon imaging, we found that HSD increased stalling of neutrophils in brain capillaries and decreased CBF. Neutrophil depletion reduced the number of stalled capillaries and restored resting CBF but did not prevent tau phosphorylation or cognitive impairment. These novel findings show that, capillary stalling contribute to CBF reduction in HSD, but not to tau phosphorylation and cognitive deficits. Therefore, the hypoperfusion caused by capillary stalling is not the main driver of the tau phosphorylation and cognitive impairment.

## INTRODUCTION

High salt intake plays a role in several human diseases and has emerged as a major public health problem worldwide^1^. Although essential for life, salt intake exceeds greatly the American Heart Association (AHA) recommended level of less than 1.5 gr/day^2^. The deleterious effects of high salt diet (HSD) have been traditionally attributed to hypertension, but accumulating evidence indicates that HSD is harmful independent of hypertension^3–7^. HSD has powerful effects on cerebral blood vessels^8–13^ and has been linked to increased incidence of cardiovascular diseases including stroke^14–22^ and dementia^23, 24^. Increasing evidence implicate HSD also in cognitive impairment^25, 26^, and lower dietary salt has been linked to improved cognitive function^27, 28^. However, the mechanisms responsible for these effects have not been fully elucidated.

In mice, we have shown that HSD reduces resting cerebral blood flow (CBF), an effect mediated by a deficit in endothelial nitric oxide (NO)^13^. However, the mechanisms by which the NO deficit leads to CBF reduction remains unclear. The NO deficit could reduce resting CBF by constricting arterioles and increasing vascular resistance^29^, but could also increase leukocyte adhesion leading to microvascular occlusions^30^. Capillary stalls, blood flow arrest by leukocytes or red blood cells (RBC) in capillaries, have emerged as a cause of CBF reduction and impaired cerebral oxygenation by increasing the heterogeneity of RBC fluxes^31, 32^, and contribute to CBF reduction in mouse models of Aβ accumulation, stroke, diabetes and seizures^33–43^. However, it is not known which of these mechanisms contribute to the hypoperfusion caused by HSD.

Another key question concerns the role of the CBF reduction in tau accumulation. We have shown that HSD-induced cognitive impairment involves the microtubule associated protein tau^44^. In tauopathies, including AD and frontotemporal dementia, tau is hyperphosphorylated, aggregates into multimers and exerts pathological effects^45^. In HSD, the endothelial NO deficit leads to de-nitrosylation of the protease calpain in neurons, which, in turn, activates the tau phosphorylating enzyme Cdk5^44^. The effect is related to lack of endothelial NO and not neuronal NO^44^. The HSD-induced cognitive impairment did not occur with administration of anti-tau antibodies or in tau^-/-^ mice, despite persistent CBF reduction^44^, indicating that tau accumulation, not hypoperfusion, is the final mediator of the cognitive dysfunction. However, the CBF reduction could also independently promote accumulation of hyperphosphorylated tau (p-Tau), as described in models of cerebral hypoxia/hypoperfusion^46, 47^, possibly via hypoxia-induced Cdk5 activation^48^.

Since capillary stalling by neutrophils has emerged as an important cause of reduced CBF in wide variety of models ^33–43^, in this study, we investigated if HSD promotes capillary stalling and whether the ensuing CBF reduction contributes to p-Tau accumulation and cognitive deficits.

## MATERIALS AND METHODS

### Mice

All procedures are approved by the institutional animal care and use committee of Weill Cornell Medicine (Animal protocol number: 0807-777A). Studies were conducted, according to the ARRIVE guidelines (https://www.nc3rs.org.uk/arrive-guidelines). To avoid the confounding effect of aging^49^, young C57BL/6 male mice obtained from Jackson Laboratory (RRID:IMSR_JAX:000664) were used.

### High salt diet

Mice (8 weeks old) received normal chow (0.5% NaCl) and tap water ad libitum (normal diet) or sodium-rich chow (8% NaCl) and tap water containing 0.9% NaCl ad libitum (HSD) for 4 to 26 weeks according to the experiment^13, 44, 50^. These levels of salt intake are an approximation of the highest levels of human salt consumption^51^.

### In vivo treatments

To selectively deplete circulating neutrophils, mice were injected intra-peritoneally (i.p.) with antibodies against the neutrophil antigen Ly6G (anti-mouse Ly6G clone 1A8, BE0075-1, Bio X Cell) or isotype IgG as a control (anti-rat IgG2a isotype control, anti-trinitrophenol, BE0089, Bio X Cell) (4mg/kg/mouse 3 times a week) for the last 4 weeks of the 12 week-HSD treatment^34, 35, 39, 42, 52^.

### Animal preparation for *In vivo* brain imaging

Optical access to brain was achieved through a thinned skull preparation, as previously described^53^. During the surgical procedure, the animals were anesthetized with isoflurane (1.5– 2%) delivered in a mixture of oxygen (75 mL/min) and nitrogen (200 mL/min) with the goal to achieve a respiratory rate of about 60 breaths per minute. To control body temperature, the animals were placed on a feedback-controlled heating blanket (TC-1000; CWE, Inc.). Following scalp and periosteum removal, a custom-made titanium head post was affixed to the right side of the parietal skull surface using a three-component metabond adhesive (S371, S398, S396, Parkell). The skull was then thinned with a dental drill, and the exposed bone surface was covered with a 3-mm diameter glass coverslip (CS-3S, Warner Instruments), using cyanoacrylate glue (Loctite 495; Henkel) to ensure increased durability. Mice were allowed to recover for at least 7 days prior to *in vivo* imaging.

### Two photon *in vivo* imaging

Mice were anesthetized with isoflurane (1-1.5%) and placed on a custom stereotactic frame. A feedback-controlled heating pad was used to control the body temperature. To label blood plasma and visualize vascular structure, 70kDa Texas Red dextran (50 µl, 2.5% w/v in saline, D-1830; Invitrogen) was injected retro-orbitally (r.o.). Leukocytes and blood platelets were labeled with Rhodamine 6G (0.1 ml, 1 mg/ml in saline, R4127; Sigma). Leukocytes were distinguished from blood platelets by Hoechst 33342 (50 μl, 4.8 mg/ml in saline, H21492; Thermo Fisher Scientific). Rhodamine 6G and Hoechst 33342 were loaded into a single syringe and r.o. injected. Isoflurane concentration was adjusted as necessary to maintain a steady breathing rate of 60 breaths/min.

Three-dimensional images of the cortical vasculature and measurement of RBCs speeds in specific vessels were acquired using a commercial 2-photon microscope (2-PEF, FVMPE; Olympus) equipped with a solid-state laser (InSight DS+; Spectraphysics). For three-channel imaging, the excitation laser was tuned to 820 nm and scanned with galvanometric scanners, with the focused beam directed into the sample using a 25× water immersion objective (XLPlan N 25L×L1.05LNA, Olympus). Emitted fluorescence passed through an infrared (IR) blocking filter (FV30-BA685RXD), was spectrally separated by a dichroic mirror (FV30-SDM570) and two filter cubes (FV30-FVG; 410-to 460-nm (Hoechst), 495-to 540-nm (Rhodamine 6G), FV30-FOCY5; 575-to 645-nm (Texas Red)). Image stacks and movies were captured using Fluoview software (FV31S-SW, version 2.3.1.163; Olympus). For vascular mapping, a low-magnification 5× objective (MPlan N 5×, 0.1 NA, Olympus) was used to acquire an initial map of the vasculature. We selected two areas with sharp contrast and avoiding large pial vessel for further imaging. The objective was then switched to 25× for higher magnification imaging. For capillary stall quantification we took high resolution stacks of images spaced by 1 µm axially to a cortical depth of 300-500 µm (1024 x 1024 resolution, frame rate of 0.31). To measure RBC speed, vessel segments parallel to the imaging surface were chosen and unidirectional line scans were performed on the line ROI drawn along the central axis inside vessel lumen for 1 minutes with a line rate of 0.8-1.3 kHz, depending on the length of line ROI. In a series of experiments comparing pre-and post-antibody treatments, the antibody was administered immediately after the initial imaging session. Two hours later, imaging was repeated at the exact same location.

### Two photon *in vivo* imaging analysis

To measure capillary segment stalling, we began by counting the number of capillary segments in each image stack. First, we created skeletons of the vascular networks using the TubeAnalyst macro (IRB Barcelona) in ImageJ. The skeletal results were then merged with the original image stack and manually corrected to address missing capillary segments due to imaging artifacts or stalling. Next, we carefully examined the image stacks, manually identifying capillary stalls by observing the movement of RBCs. Since Texas Red dextran labels blood plasma and not the blood cells, RBCs appear as dark patches or stripes in the vessel lumen. The consistent presence of these dark stripes indicates ongoing blood flow in that vessel segment. We plotted the stall counts as the fraction of capillaries with stalled blood flow. Stalls were further categorized based on our labeling strategy: capillary segments with a cell-shaped object labeled with both Rhodamine 6G and Hoechst were considered to contain a leukocyte, while those with punctate Rhodamine 6G labeling alone were categorized as platelet aggregates (Fig. 1A). Stalled segments with only RBCs present were classified as RBC stalls. Additionally, we categorized stalls based on their duration: those that persisted throughout the imaging were labeled as “stalled,” while those that resolved and resumed blood flow were classified as “transient.” Each frame captured took 3.2 seconds, with capillary diameters typically ranging from 6 to 9 µm, and stacks were taken at 1 µm intervals. As a result, each capillary remained visible for approximately 28.8 seconds.

**Figure 1.**
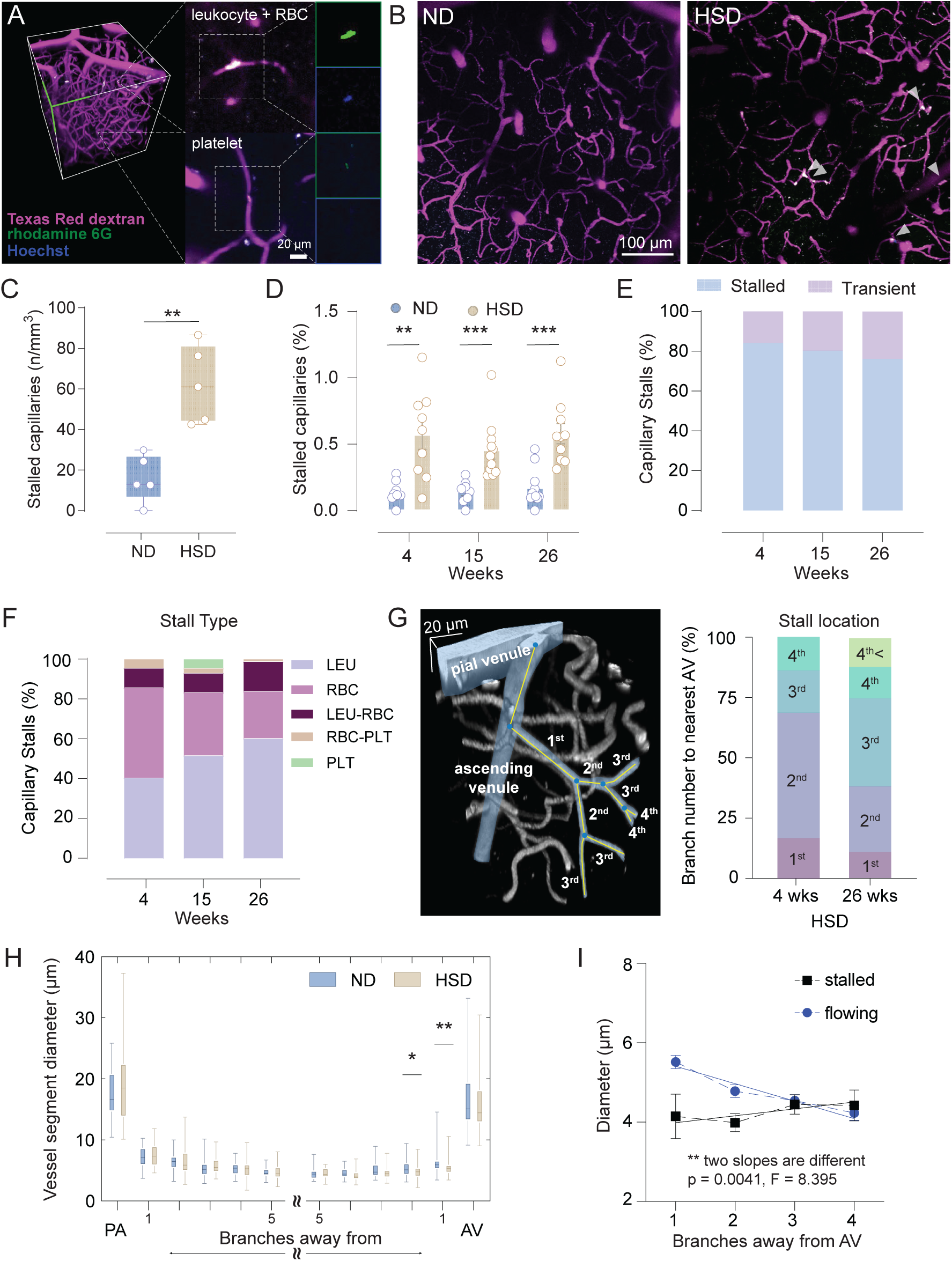
**HSD increases brain capillary stalling**. **A**. Rendering of *in vivo* 2PEF image stack of cortical vasculature (left), and images of stalled capillaries that contained a leukocyte and RBC (right, top), and platelet (right, bottom), distinguished by fluorescent labels (magenta: Texas Red dextran labeled blood plasma; green: rhodamine 6G labeled leukocytes and platelets; blue: Hoeschst labeled leukocyte nuclei). **B**. Representative 100 µm axial projections of 2PEF image stacks from ND (left) and HSD (right) diet fed mice showing stalled capillaries indicated with arrowheads. **C**. HSD (15wks) increases the number of stalled capillaries per volume of tissue (p=0.0017 vs ND, n=5/group, two-tailed paired t-test). **D**. Fraction of capillaries with stalled blood flow in ND and HSD fed mice over time. Each data point represents fraction of capillaries stalled in one stack. (Diet: p<0.0001, Time: p=0.4855, Interaction: p=0.6126, two-way ANOVA plus Tukey’s multiple comparisons test; 4 weeks: p=0.0002 vs ND, 15 weeks: p=0.0188 vs ND, 26 weeks: p=0.0003 vs ND) **E**. Fraction of stalled capillaries that remained stalled (light blue) or resumed flowing during stack acquisition (light violet) in HSD mice over time (Weeks of HSD: p>0.9999, stalls time: p<0.0001, Interaction: p=0.5806, n=5/group, two-way ANOVA plus Tukey’s multiple comparisons test). **F.** Fraction of stalled capillaries in HSD mice that contained leukocytes (LEU), one or more RBCs and platelets (PLT), distinguished as LEU only (LEU), LEU with one or more RBCs (LEU-RBC), RBCs with PLT (RBC-PLT) and PLT only (Weeks of HSD: p>0.9999, stalls type: p<0.0001, Interaction: p=0.1880, n=5/group, two-way ANOVA plus Tukey’s multiple comparisons test). **G**. Vessel enumeration scheme overlay onto rendering of an ascending venule and its branches (left) and stall location in each capillary branch (right) in HSD mice (4 and 26 weeks). **H.** Diameter of randomly chosen branches from PA and AV (20-76 vessel segments for each column from ND and HSD fed mice, 1^st^ branch from AV: p=0.0049, 2^nd^ branch from AV: p=0.0378, n=5/group, two-tailed Mann-Whitney tests). **I**. Diameter from 1^st^ to 4^th^ branch from AV shown in **H** broken down by the presence of capillary stall. Analysis of covariance (ANCOVA) test to compare two slopes. Data is presented as mean ± SEM.

Blood vessel segments were categorized as either penetrating arterioles (PA), capillaries, or ascending venules (AV). Arterioles were differentiated from venules based on their morphology—arterioles had smaller diameters, smoother walls, and more symmetrical Y-shaped branching compared to venules. Smaller branch segments were classified by tracing their connections to these easily recognizable larger vessels. For each identified stall, we traced the upstream and downstream branches and examined the proximity to either PA or AV. While one stall was found at the 4^th^ branch near the PA, most stalls were located near the AV. We quantified the branching order relative to the AV, following the vessel enumeration scheme shown in Fig. 1F.

Capillary diameter was measured using ImageJ (NIH). For each high-resolution stack, a projection stack was generated, with each frame consisted of maximum z-projection of every 10 frames, using a custom-written ImageJ macro. Per each projection stack, 50 vessels were randomly selected, ensuring representation across different vessel categories, spanning from PA and its 1st to 5th branches and 5th to 1st branches toward AV and AV. To select the most appropriate frame for each vessel, we scrolled through the z-axis of the projection stack and identified the frame where the vessel of interest was most visually prominent. We draw a perpendicular line across the vessel segment, and the full width at half maximum (FWHM) was measured using the “Plot Profile” tool in ImageJ.

Space-time images were generated from line scans, which exhibited diagonal streaks. The slope of these streaks was inversely proportional to the centerline RBCs speed. All line scans were analyzed using MATLAB, which employed a Radon transform-based algorithm to compute line integrals of the 2D image [g(x,y)] at various angles (https://github.com/sn-lab/Blood-flow)^54, 55^. To estimate the RBC speed of each vessel segment, we averaged the RBCs speed measurements over a 1-minute period. Volumetric blood flow (*F*) was then calculated using the following equation: *F=πvr^2^/2* where *v* is the centerline RBC speed derived from line scan and *r* is the vessel radius. All displayed 2PEF images were generated using ImageJ (NIH) and represent maximal projections of 2PEF image stacks except for the renderings in Fig. 1, which utilized 3D viewer tool.

### Laser speckle imaging and analysis

Laser speckle imaging (LSI) was performed with a commercial system (Omegazone, Omegawave). Cranial window was illuminated with a semiconductor laser set at 780 nm, and the scattered light was detected with a CCD camera positioned over the skull. Animals were anesthetized with isoflurane (1∼1.5%), and after a 10-minute of stable baseline, they were administered either anti-Ly6G or isotype control antibodies (4 mg/kg, i.p). Since neutrophil depletion from a single injection of anti-Ly6G lasted for 1 week (data not shown), we gave alternating treatments to same group of animals 2 weeks apart. The mice were then monitored and imaged for additional two hours. Color-coded CBF images were obtained at the highest resolution (639L×L480 pixels) with the sampling frequency of 1 Hz. To obtain a single representative frame for each minute of measurement, 60 consecutive frames were averaged, resulting in a time series of averaged frames. To account for spatial drift across the time series, we used the ‘rigid body transformation’ using the ImageJ (NIH) TurboReg plugin. Speckle contrast values were acquired from whole cranial window, summing all contrast from surface arteriole, surface venule, or parenchymal region. This time-series data was subsequently analyzed for plotting and quantification at the 2-hour time point.

### Isolation and flow cytometry analysis of peripheral leukocytes

For isolation of peripheral leukocytes, mice were anesthetized with pentobarbital (100 mg kg–1, i.p) and 0.5 ml of blood was collected by cardiac puncture into heparinized tubes. Blood (150 µL) was incubated for 5 minutes at room temperature with 5 mL of erythrocytes lysis buffer followed by addition of 10 mL of HBSS/10mM HEPES and spun at 500g for 7 minutes. Afterward, blood cells were resuspended in MACS buffer (PBS supplemented with 2% FBS, 2 mM EDTA). Cells were resuspended in 50 µL of FACS buffer (PBS with 2% FBS and 0.05% NaN_3_), blocked with anti-CD16/CD32 for 10 minutes at 4°C and then stained with the appropriate antibodies for 15 minutes at 4°C. The following antibodies were used: CD45 (clone 30F-11), Ly6C (clone HK1.4), Ly6G (clone 1A8), Cxcr4 (clone L276F12), Gr-1 (clone RB6-8C5), CD182 (clone SA044G4), CD11b (clone M1/708) from Biolegend.

### Isolation and flow cytometry analysis of brain cells

For isolation of brain cells, mice were anesthetized with pentobarbital (100 mg kg–1, i.p.) and transcardially perfused with heparinized PBS. Brain hemispheres were separated from the cerebellum and olfactory bulbs and gently triturated in HEPES-HBSS buffer (138 mM NaCl, 5 mM KCl, 0.4 mM Na_2_HPO_4_, 0.4 mM KH_2_PO_4_, 5 mM D-Glucose, 10 mM HEPES) using a Gentle MACS dissociator (Miltenyi Biotec) following the manufacturer’s instructions. The suspension was digested with 62.5 µg/ml Liberase DH (Roche Diagnostics), 0.8 U/ml dispase (Worthington) and 50 U/ml DNAse I (Worthington) at 37°C for 45 min in an orbital shaker at 100 rpm. Isolated cells were washed and centrifuged in 30% Percoll (GE Healthcare) for myelin removal. Cells were resuspended in 50 µL of FACS buffer (PBS with 2% FBS and 0.05% NaN_3_), blocked with anti-CD16/CD32 for 10 minutes at 4°C and then stained with the appropriate antibodies for 15 minutes at 4°C. The following antibodies were used: CD45 (clone 30F-11), CD11b (clone M1/70), Ly6G (clone 1A8), Ly6C (clone HK1.4), CD62L (clone MEL-14), ICAM/CD54 (clone YN1/1.7.4), CD18 (clone M18/2) from Biolegend^13, 56^, CD62E (clone 10E9.6) from BD Pharmingen and CD62P (clone Psel.KO2.3) from Thermo Fisher. For both brain and blood flow cytometry, cells were washed with FACS buffer, resuspended in 150 µL of FACS buffer and acquired in a NovoSampler Q (NovoCyte Quanteon). Appropriate isotype controls, “fluorescence minus one” staining, and staining of negative populations were used to establish sorting parameters. Analysis was performed with FlowJo.

### Positive selection of circulating neutrophils

Blood neutrophils were purified by immunomagnetic positive selection. Briefly, 500 µL of blood was withdrawn from the heart using a 0.5M-EDTA flushed syringe. Two rounds of erythrolysis were performed as described for flow cytometry analysis of peripheral leukocytes. Leukocytes were resuspended in MACS buffer (PBS supplemented with 2% FBS, 2 mM EDTA; 300µl/10^7^ cells), incubated with anti-Ly6G biotinylated-antibody (750ng/10^7^ cells; clone 1A8; # 127604, Biolegend) and purified with anti-biotin-microbeads according to the manufacturer instructions (Miltenyi Biotech). Afterwards, cells were washed, centrifuge at 1500, 7min, 4°C, and resuspended in 1ml Trizol reagent (Invitrogen Life Technologies).

### Quantitative Real-Time PCR

Quantitative determination of gene expression was examined as described before^13^. RNeasy Plus Mini Kit (Qiagen) was used to extract RNA from circulating neutrophils. After, cDNA was synthetized with iScript reverse transcription supermix for RT-qPCR (BioRad). qRT-PCR was conducted with cDNA in duplicate reactions using the Maxima SYBR Green/ROX qPCR Master Mix (2X) (Thermo Scientific). The reactions were incubated at 50°C for 2 minutes and then at 95°C for 10 minutes. A polymerase chain reaction cycling protocol consisting of 15 seconds at 95°C and 1 minute at 60°C for 45 cycles was used for quantification. Primer sequences (Invitrogen Life Technologies) are described in Table 1. Data were expressed as relative fold change over ND mice calculated by the 2^-ΔΔCt^ method^57^.

### Immunoblot analysis of phosphorylated tau

Cortex (≈90-100mg) and hippocampus (≈15-20mg) isolated from ND and HSD mice were sonicated in 800 and 500μl of RIPA buffer (50mM Tris-HCl pH 8.0, 150mM NaCl, 0.5% Deoxycholic Acid, 0.1% SDS, 1mM EDTA pH 8.0, 1% IGEPAL CA-630, 1mM Na3VO4, 20mM NaF and one tablet/10mL of cOmplete™, EDTA-free Protease Inhibitor Cocktail, Millipore Sigma). After homogenization in cold RIPA buffer and centrifugation (20.000*g* for 10 minutes), 150µl of the supernatant containing the proteins was boiled at 100°C for 10 minutes. Samples were cooled on ice for 20 minutes and then centrifuged at 20,000 g at 4°C for 15 minutes. The supernatant corresponding to the heat stable (HS) fraction was then harvested. The proteins were then mixed with equal volumes of SDS sample buffer, boiled, and analyzed on 10% Novex™ WedgeWell™ gels (Thermo Fisher). Proteins were transferred to PVDF membranes (Millipore), blocked at room temperature (RT) for 1 hour with 5% milk in TBS, and incubated, overnight at 4°C, with primary antibodies (see Table 2) in 5% BSA in TBS/0.1% Tween-20 (TBST). Membranes were washed in TBST, incubated with goat anti-mouse or rabbit secondary antibodies conjugated to horseradish peroxidase (Santa Cruz Biotechnology) for 1 hour at RT and protein bands were visualized with Clarity Western ECL Substrate (Bio Rad) on an iBright™ CL1500 Imaging System (Thermo Fisher). Quantification was performed using iBright Analysis Software (Thermo Fisher).

### Barnes Maze test

The Barnes maze consisted of a circular open surface (90cm in diameter) elevated to 90cm by four wooden legs. There were 20 circular holes (5cm in diameter) equally spaced around the perimeter and positioned 2.5cm from the edge of the maze. No wall and no intra-maze visual cues were placed around the edge. A wooden plastic escape box (11×6×5cm) was positioned beneath one of the holes. Three neon lamps and a buzzer were used as aversive stimuli. The Any-Maze tracking system (Stoelting) was used to record the movement of mice on the maze. Extra-maze visual cues consisted of objects within the room (table, computer, sink, door, etc.) and the experimenter. Mice were tested in groups of ten, and between trials they were placed into cages, which were placed in a dark room adjacent to the test room for the inter-trial interval (20-30 minutes). No habituation trial was performed. The acquisition phase consisted of 3 consecutive training days with three trials per day with the escape hole located at the same location across trials and days. On each trial a mouse was placed into a start tube located in the center of the maze, the start tube was raised, and the buzzer was turned on until the mouse entered the escape hole. After entering the escape box, mice were returned to their cage.

Between trials the maze floor was cleaned with 10% ethanol in water to minimize olfactory cues. For each trial mice were given 3 minutes to locate the escape hole, after which they were guided to the escape hole or placed directly into the escape box if they failed to enter the escape hole. Four parameters of learning performance were recorded: (1) the latency to locate (primary latency) and (2) enter the escape hole (total latency), (3) the number of errors made and (4) the distance traveled before locating the escape hole. When a mouse dipped its head into a hole that did not provide escape was considered an error. On day 4, the location of the escape hole was moved 180° from its previous location (reverse learning) and two trials were performed^13, 44^.

### Nest building

The ability of mice to build nests was assessed by the Deacon rating scale^58^. One hour before the dark cycle, each mouse was placed in a new clean cage containing 2.5-3.0 g of nestlet (Ancare). Food, water and lighting parameters were not changed from standard housing practices. The next day, nests were assessed on a rating scale of 1–5. and untorn nestlet pieces were weighed. The cognitive parameters recorded were (1) nest score and (2) percentage of untorn nestlet^13^.

### Statistical analysis

Sample size was determined according to power analysis based on previously published work by our lab on the effects of dietary salt on CBF regulation and cognitive function^13,44^. On these bases, 10-15 mice/group were required in studies involving assessment of cognitive function^13,59^. Mouse randomization was performed based on the random number generator function (RANDBETWEEN) in Microsoft Excel software. After testing for normality (Shapiro-Wilk, D’Agostino or Pearson test), intergroup differences were analyzed by unpaired two-tailed t-test for single comparison, or by one-way ANOVA or two-way ANOVA with Tukey’s, Bonferroni’s, or Sidak’s multiple comparison test, as appropriate and indicated in the figure legends. If non-parametric testing was indicated, intergroup differences were analyzed by Mann-Whitney test, Wilcoxon matched-pairs test or Kruskal-Wallis test with Dunn’s correction, as appropriate and indicated in the figure legends. Statistical tests through the manuscript were performed using Prism 10 (GraphPad). Data are expressed as mean±SEM and differences are considered statistically significant for p<0.05. Statistical significance is represented on the plots as follows: *p< 0.05; **p< 0.01; ***p< 0.001.

## RESULTS

### High salt diet promotes capillary stalling

To examine whether HSD increases capillary stalling, we used *in vivo* 2PEF to image the cortical vasculature in C57BL6/J male mice fed a ND or HSD from 4 to 26 weeks (Fig. 1A-B). Capillary segments present in the stacks were individually assessed to be either flowing or stalled based on RBCs motion while the capillary is visible during three-dimensional image stack acquisition for several seconds (Fig. 1A-B). Since Texas Red dextran was used to label plasma, blood cells appear as dark moving intraluminal patches. A capillary segment was scored as stalled if there was no motion of the RBCs and other cells in the Texas Red imaging channel (Fig. 1A-B). *In vivo* fluorescent labeling by i.v. administration of rhodamine 6G and Hoechst was used to label blood leukocytes (Fig.1A). We found that the number of stalled capillaries, expressed as percentage of total capillaries present in the stack, was increased in HSD mice (Fig. 1B-C). In agreement with previous studies^34, 40^, our results indicate that around 0.5 % of capillaries in HSD mice had stalled blood flow, while in age matched ND mice around 0.1 % of capillaries were not flowing (Fig. 1C). In HSD mice, capillary stalling was elevated starting at 4 weeks and remained higher than ND mice up to 26 weeks (Fig. 1D). Most stalled capillaries were occluded for the entire duration of the stack acquisition (Fig. 1E). However, we found the about 20% of stalls disappeared during stack acquisition, suggesting transient stalling (Fig. 1E). By using rhodamine 6G and Hoechst to identify blood leukocytes, we found that most stalled capillary segments contained leukocytes and/or red blood cells (Fig. 1E). Next, we assessed the location of the stalls based on the order of the capillary branches nearest to the PA or AV (Fig. 1F)^60^. At both 4 and 26 weeks, we found that almost all stalls were found near AV, in the 2^nd^ and 3^rd^ branches from AV (Fig.1F), consistent with the venules being more prone to leukocyte adhesion particularly in conditions, like HSD, associated with NO deficit^61^. Next, we randomly selected capillaries in cortical stacks and measured their diameter. Consistent with the location of the stalls (Fig. 1F), we found that branches closer to AV (1^st^ and 2^nd^ branches) had smaller diameter (Fig. 1G-H). Furthermore, we found that the diameter of stalled capillaries was significantly reduced compared to that of flowing capillaries (Fig. 1H).

### Neutrophil depletion reduces the number of stalled capillaries and increases CBF

We then examined the effect of capillary stalling on CBF. To reduce the number of capillary stalling, we depleted neutrophils by administering anti-Ly6G antibodies^34, 62^. Anti-Ly6G treatment removed approximately 58% of the capillary stalls caused by HSD (Fig. 2A). Next, to assess the impact of anti-Ly6G administration on brain perfusion we used LSI. We observed that anti-Ly6G lead to a ∼15% increase in neocortical perfusion compared to baseline in HSD mice (Fig. 2B-D). To further characterize the effects of neutrophil depletion on brain perfusion at the microcirculation level, we used 2PEF. RBC speed was measured by acquiring line-scan images on capillary segments before and after i.p. administration of anti-Ly6G or IgG in ND and HSD mice. We found that anti-Ly6G administration, increased RBC speed in PA and its capillary branches (Fig. 2E). Additionally, we found that neutrophil depletion increases the diameter of the 1^st^ branch closer to AV (2F). Finally, the volumetric blood flow, which accounts for both RBC speed and vessel diameter, was increased in both the PA and AV (Fig. 2F). Overall, these data indicate that neutrophil depletion reduces the fraction of stalled capillaries and increases CBF in both the PA and the AV.

**Figure 2.**
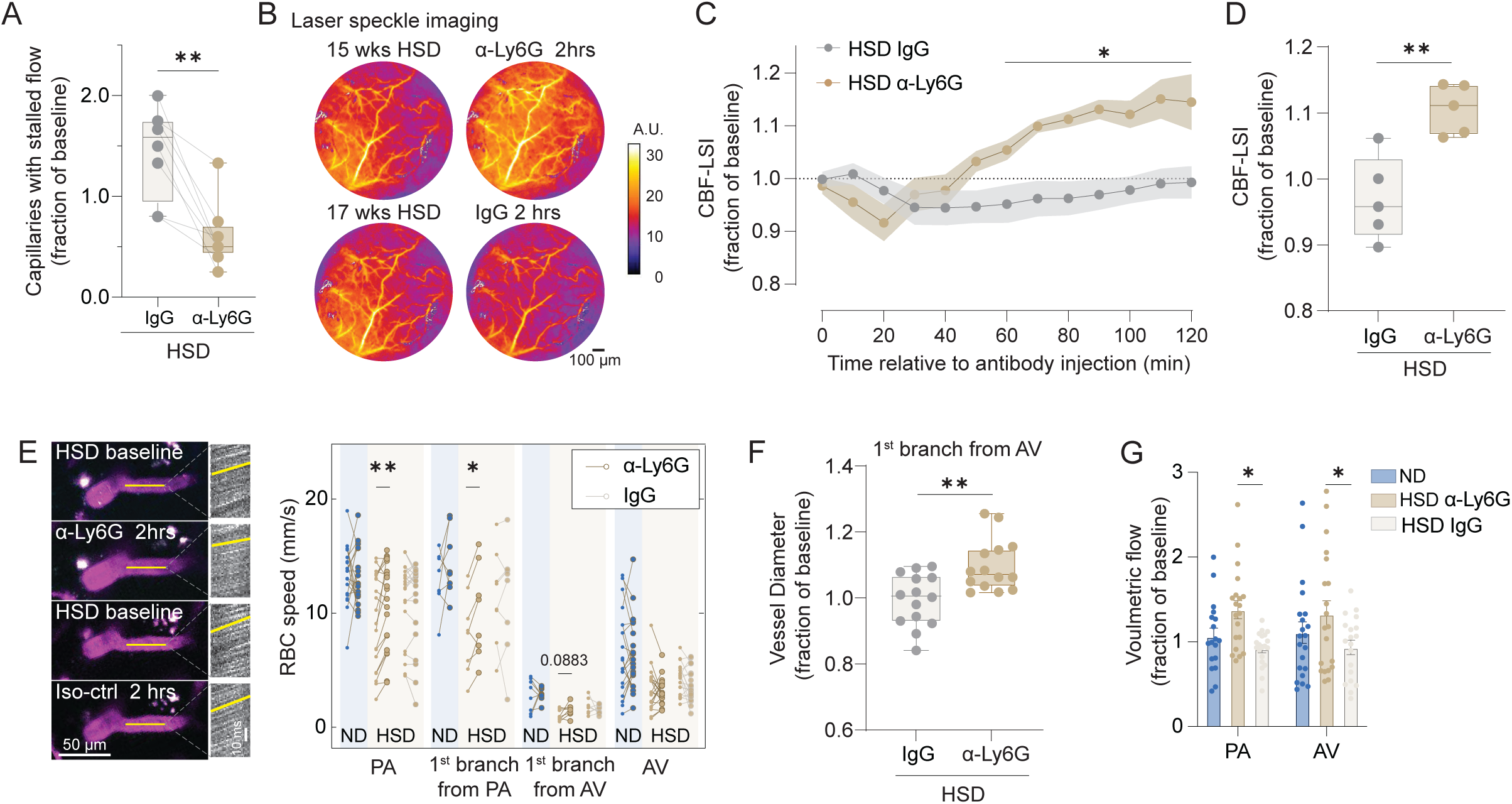
Neutrophil depletion rescues CBF in HSD fed mice. **A**. Fraction of capillaries with stalled blood flow 2 hrs after anti-Ly6G (α-Ly6G) or Iso-ctrl (IgG) antibody administration in HSD fed mice (p=0.0078, n = 5/group, Wilcoxon matched-pairs signed rank test). **B**. Representative LSI images of HSD mice injected either with IgG or α-Ly6G. **C**. LSI CBF plot as a function of time after antibody injection (Time: p<0.0001, Treatment: p=0.0337, Interaction: p<0.001, n=5/group, two-way ANOVA plus Sidak’s multiple comparisons test). **D**. Average CBF-LSI increase from 60 to 120 min after IgG/ α-Ly6G injection (p=0.0034 vs HSD IgG, n=5/group, two-tailed unpaired t-test). **E**. Representative axial projection of 1^st^ branch from an AV of HSD mouse, before and 2 hrs after alternating antibody treatments. Corresponding line scan (left) and plot of RBCs speed (right) before and 2 hrs after antibody treatment (HSD α-Ly6G PA: p=0.0030, n=20 from 5 mice, HSD α-Ly6G 1^st^ branch from PA: p=0.0104, n=8 from 5 mice, two-tailed paired t-test). **F**. Relative vessel diameter of 1^st^ branch from AV after IgG or α-Ly6G injection (p=0.0019 vs HSD IgG, n=5/group, two-tailed unpaired t-test). **G.** Volumetric blood flow in PA and AV 2 hrs after α-Ly6G or IgG administration in ND and HSD mice shown as a fraction of baseline flow (PA: p=0.0101, n=20 from 5 mice, AV: p=0.0378, n=20 from 5 mice, Tukey’s multiple comparisons test). Data is presented as mean ± SEM.

### HSD does not induce activation of circulating or brain neutrophils

To investigate whether the increase in capillary stalling in HSD could be attributed to neutrophil activation, which in turn could promote adhesion to brain endothelial cells (ECs), we performed blood and brain flow cytometry in ND and HSD mice. We found that the number and the frequency of blood and brain neutrophils (CD45^hi^CD11b^+^Ly6G^+^), increased slightly in HSD, albeit not significantly (Fig. 3B and 4B). Circulating numbers of other myeloid sub-populations (CD45^hi^CD11b^+^Ly6C^hi^ and CD45^hi^CD11b^+^Ly6C^low^) were also not significantly altered in HSD mice (Fig. 3C). Next, we assessed expression of markers of neutrophil activation including CD18, a component of LFA1 (CD11a/CD18) and MAC1 (CD11b/CD18) and L-selectin (CD62L), the only selectin expressed on neutrophils^63, 64^. We found that, compared to mice on ND, the expression of both molecules was not changed in blood and brain neutrophils isolated from HSD mice (Fig. 3D and 4C). Similarly, the expression of mediators associated with neutrophil activation (*Cxcl2*, *Cybb*, *Mmp9* and *Il1b)*^65^, assessed by RT-PCR, was not altered in circulating neutrophils isolated from HSD mice (Fig. 3E). Finally, consistent with previous data^13^, we found that expression of adhesion molecules [CD54 (ICAM), CD62E (E-selectin) and CD62P (P-Selectin)] in brain ECs was similar in ND and HSD mice (Fig. 4D). Therefore, these findings suggest that HSD does not induce activation of brain or circulating neutrophils and brain ECs.

**Figure 3.**
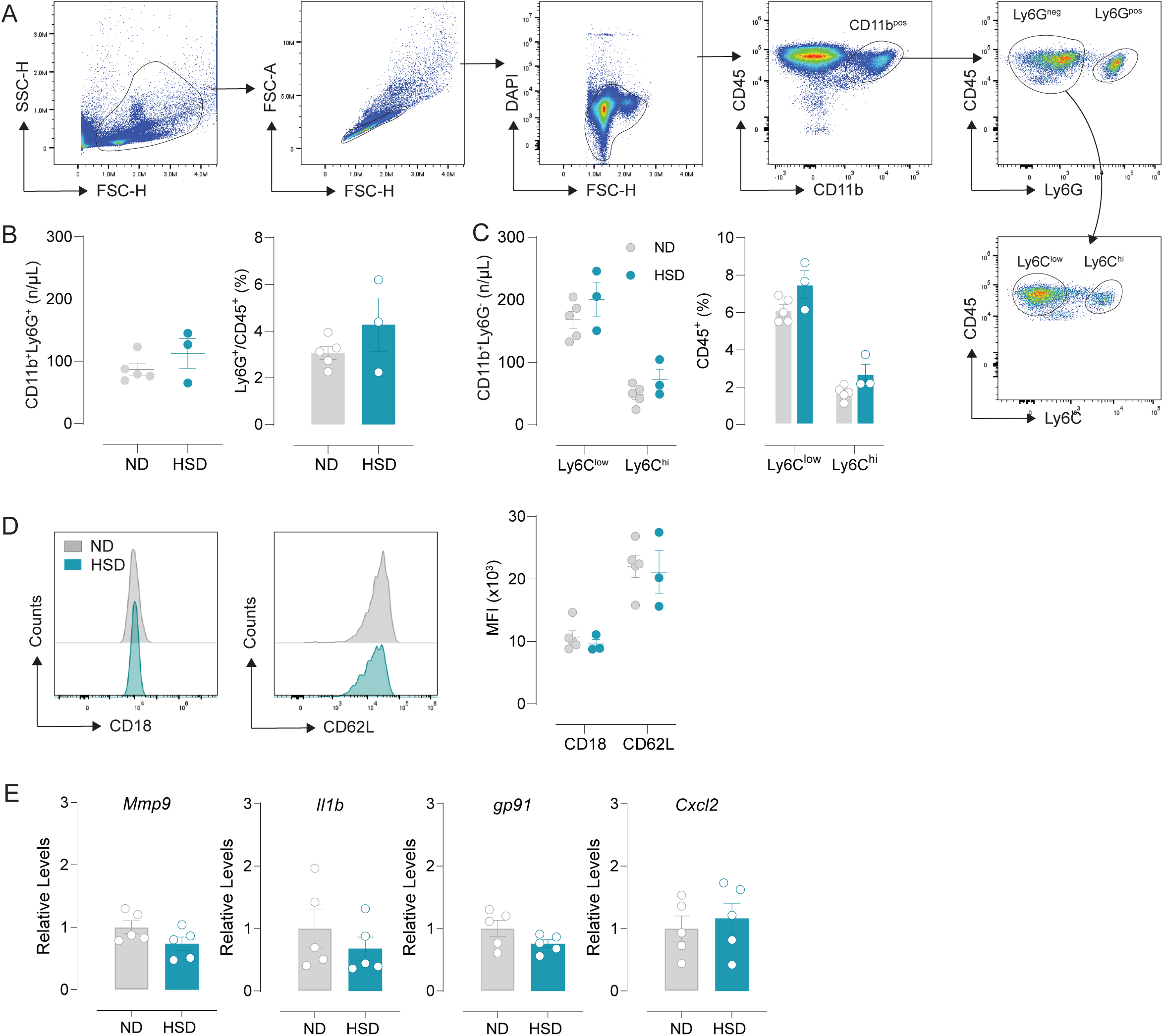
HSD does not induce blood neutrophil activation. **A**. Representative flow cytometry plots showing the gating strategy used to identify blood neutrophils (CD45^+^CD11b^+^Ly6G^+^). **B**. HSD had no effect on the number and the frequency of blood neutrophils (Numbers: p=0.2902 vs ND; Frequency: p=0.2347 vs ND, n=3-4/group, two-tailed unpaired t-test). **C**. HSD did not alter the number and the frequency of blood monocytes (Ly6C^low^, Numbers: p=0.2734 vs ND; Frequency: p=0.0829 vs ND; Ly6C^hi^, Numbers: p=0.1705 vs ND; Frequency: p=0.0603 vs ND n=3-4/group, two-tailed unpaired t-test). **D**. Representative flow cytometry plot showing CD18 and CD62L median fluorescence intensity (MFI) in blood neutrophils. HSD does not alter MFI levels of CD18 and L-selectin in blood neutrophils (CD18: p=0.4751 vs ND; L-selectin: p=0.7964 vs ND, n=3-5/group, two-tailed unpaired t-test). **E**. HSD does not increase mRNA levels of neutrophils activation markers in circulating neutrophils (*Mmp9*: p=0.1203 vs ND; *Il1b*: p=0.3884 vs ND; *Gp91*: p=0.1355 vs ND; *Cxcl2*: p=0.6248 vs ND, n=5/group, two-tailed unpaired t-test). Data is presented as mean ± SEM.

**Figure 4.**
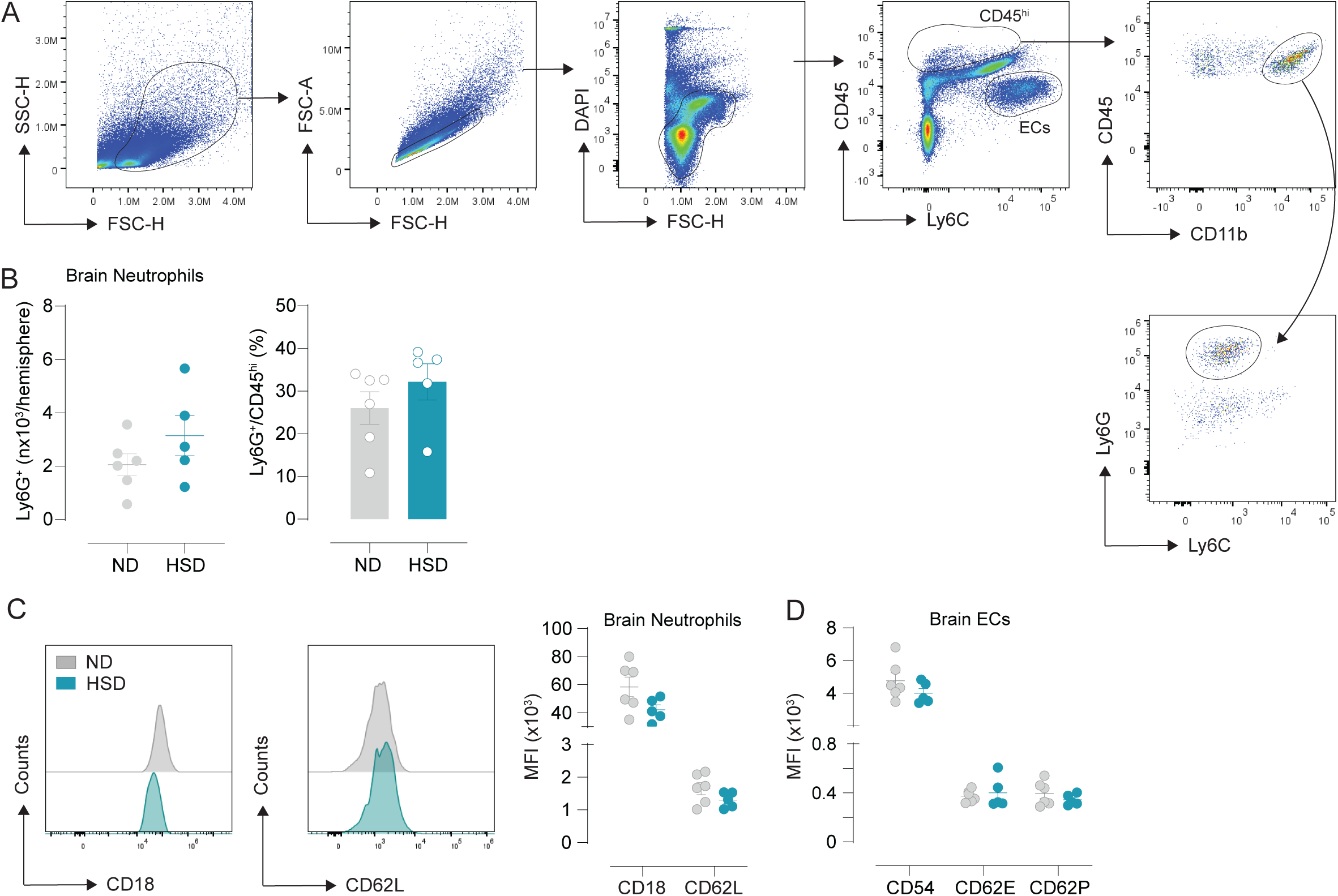
HSD does not induce brain neutrophil activation. **A**. Representative flow cytometry plots showing the gating strategy used to identify brain neutrophils (CD45^hi^CD11b^+^Ly6G^+^). **B**. HSD increases, albeit not significantly, the number and the frequency of brain neutrophils (Numbers: p=0.2154 vs ND; Frequency: p=0.3084 vs ND, n=5-6/group, two-tailed unpaired t-test). **C**. Representative flow cytometry histograms showing CD18 and CD62L median fluorescence intensity (MFI) in brain neutrophils. HSD does not alter MFI levels of CD18 and CD62L in brain neutrophils (CD18: p=0.0826 vs ND; CD62L: p=0.1582 vs ND, n=5-6/group, two-tailed unpaired t-test). **D**. HSD does not increase CD54, CD62E and CD62P median fluorescence intensity (MFI) in brain ECs (CD54: p=0.2326 vs ND; CD62E: p=0.6569 vs ND; CD62P: p=0.3108 vs ND, n=5-6/group, two-tailed unpaired t-test). Data is presented as mean ± SEM.

### Neutrophil depletion does not rescue the cognitive impairment associated with HSD

Next, we examined the effects of neutrophil depletion on the cognitive impairment associated with HSD. Mice were injected with either antibodies against Ly6G or isotype IgG as a control (4mg/kg/mouse 3x week; i.p.) for the last 4 weeks of the 12 weeks ND/HSD treatment. Cognitive function was first assessed by the Barnes maze test^13^. We found that ND and HSD mice, injected either with IgG isotype control or anti-Ly6G antibodies, were able to learn the location of the escape hole over the 3-days training phase (Fig.5A). Consistent with previous findings, when the escape hole was moved to the opposite quadrant on day 4, HSD mice spent more time to find the novel location that of ND mice^13, 44^. Importantly, we found that injection of either IgG isotype control or anti-Ly6G antibodies, had no effect on the cognitive impairment of HSD mice (Fig.5B).

**Figure 5.**
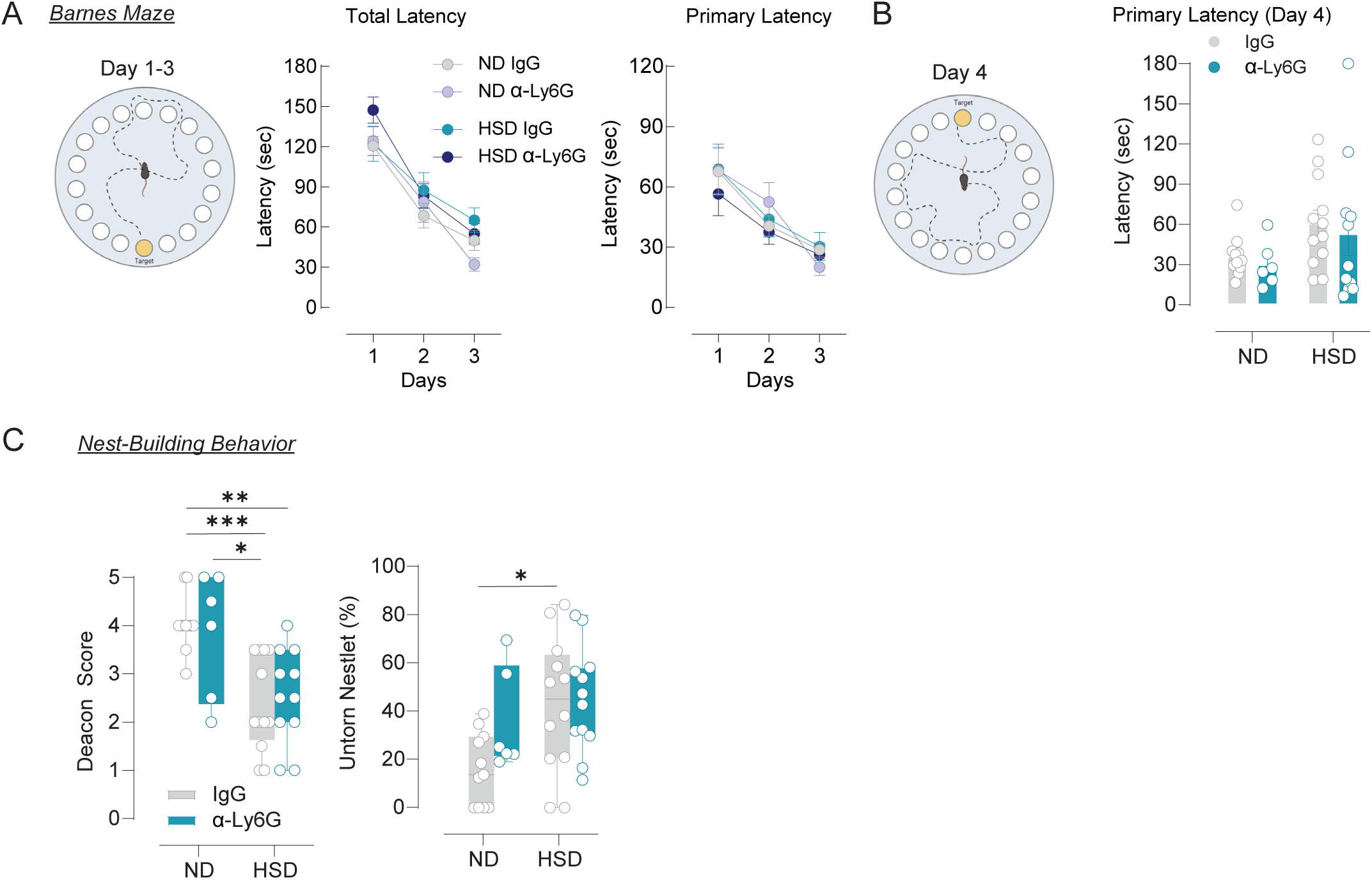
Neutrophil depletion does not rescue the cognitive impairment associated with HSD. **A.** Injection of IgG control or anti-Ly6G antibodies does not affect the learning process of ND and HSD mice during the 3-days training phase (Total latency, Time: p<0.0001, Diet: p=0.1062, Treatment: p=0.7963; Primary latency, Time: p<0.0001, Diet: p=0.7053, Treatment: p=0.6101, n=7-12/group three-way ANOVA plus Tukey’s multiple comparisons test). **B.** Primary latency is increased in HSD mice, regardless of the injection of IgG control or anti-Ly6G antibodies, on the 4^th^ day of testing when the location of the escape hole is moved 180° from its previous location (Diet: p=0.0487, Treatment: p=0.5498, Interaction: p=0.9458, n=7-12/group, two-way ANOVA plus Tukey’s multiple comparisons test). **C.** Nest building ability is impaired in both IgG control and anti-Ly6G antibodies-injected HSD mice. The amount of untorn nestlet is also increased in both IgG control and anti-Ly6G antibodies injected HSD mice (Deacon Score, Diet: p<0.0001, Treatment: p=0.9466, Interaction: p=0.5398; Untorn nestlet, Diet: p=0.0196, Treatment: p=0.1339, Interaction: p=0.2454, n=6-12/group, two-way ANOVA plus Tukey’s multiple comparisons test). Data is presented as mean ± SEM.

We also examined whether neutrophil depletion rescues the impairment of nesting behavior observed in HSD mice^13^. We assessed the quality of the nest by the Deacon score, and we measured the amount of nesting material used. We found that HSD disrupted the ability of mice to build a nest as indicated by scores between 1 and 4 and by the increase in the percentage of unshredded nestlet (Fig.5C). However, as with the Barnes maze, depletion of neutrophils had no effect on the impairment of nest building behavior of HSD mice (Fig. 5C). After completion of cognitive testing, depletion of circulating neutrophils was validated by flow cytometry (Fig. 6A-B). Taken together, these data indicate that neutrophil depletion has no effects on the cognitive impairment of HSD mice.

**Figure 6.**
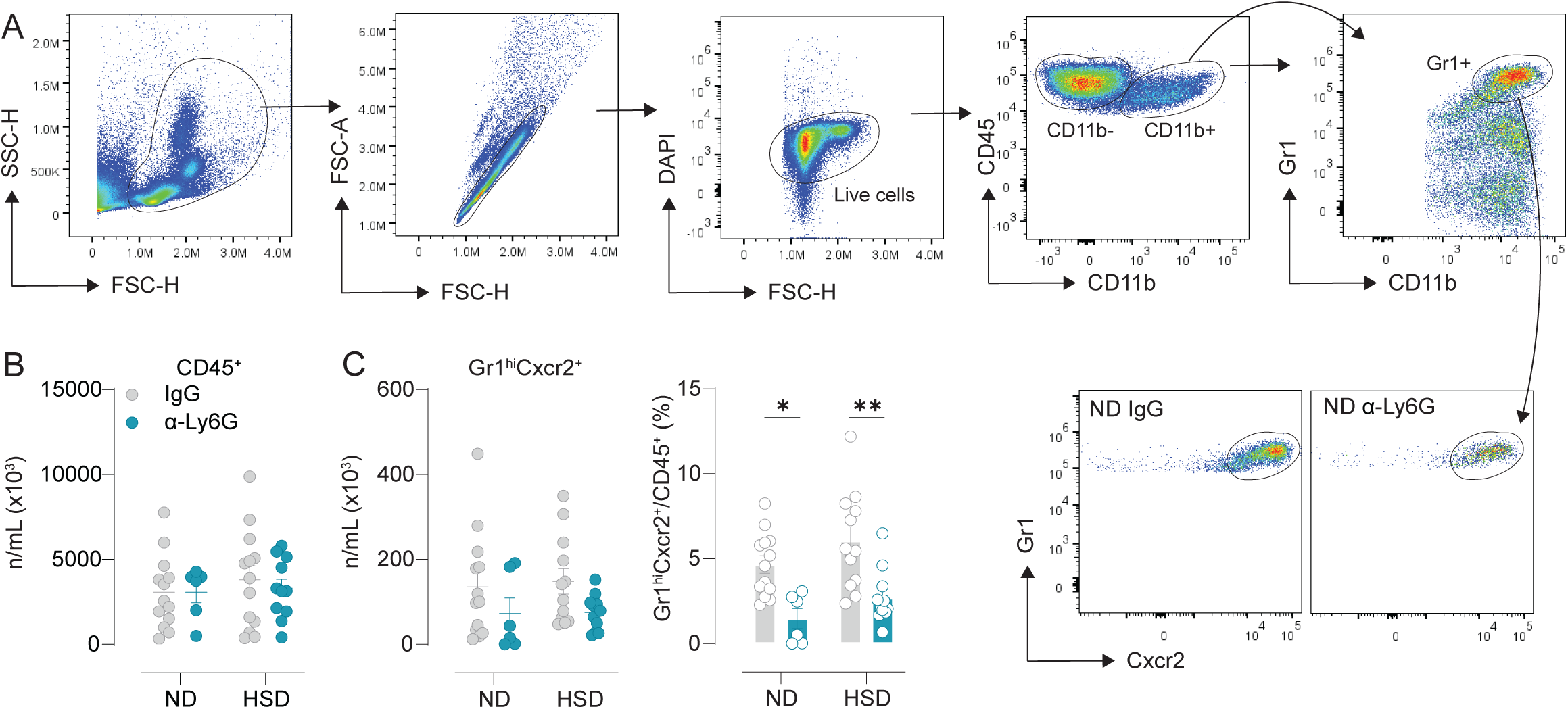
Administration of anti-Ly6G antibodies leads to depletion of circulating neutrophils in ND and HSD mice. **A.** Representative flow cytometry plots showing the gating strategy used to identify blood neutrophils (CD45^+^CD11b^+^Gr1^+^Cxcr2^+^). **B.** Injection of IgG control or anti-Ly6G antibodies does not affect the number of blood leukocytes (DAPI^neg^CD45^+^) in both ND and HSD mice (Diet: p=0.5102, Treatment: p=0.7347, Interaction: p=0.7256, n=6-13/group, two-way ANOVA plus Tukey’s multiple comparisons test). **C.** Injection of IgG control or anti-Ly6G antibodies reduces the numbers and frequency of blood leukocytes (DAPInegCD45+) in both ND and HSD mice (Numbers, Diet: p=0.8088, Treatment: p=0.0388, Interaction: p=0.8729; Frequency, Diet: p=0.0634, Treatment: p<0.0001, Interaction: p=0.9156, n=6-13/group, two-way ANOVA plus Tukey’s multiple comparisons test). Data is presented as mean ± SEM.

### Neutrophil depletion does not suppress the increase in phosphorylated tau induced by HSD

Since the cognitive impairment associated with HSD is dependent on tau accumulation^44^, we next investigated whether neutrophil depletion has any effect on the tau phosphorylation increase induced by HSD. Consistent with previous data^44^, we found that HSD increased tau phosphorylation on Ser^202^Thr^205^ (AT8) and Thr^231^ in both cortex and hippocampus of mice injected with IgG isotype control antibodies (Fig. 7A-B). We also found that phosphorylation of Thr^217^ was increased in the cortex but not in the hippocampus of the same group of mice (Fig. 7A-B). However, in agreement with the behavioral data, we found treatment with anti-Ly6G antibodies failed to attenuate phosphorylated tau levels in HSD mice (Fig. 7A-B). Thus, the data indicate that neutrophil depletion has no effect on the tau phosphorylation increase observed in the brains of HSD mice.

**Figure 7.**
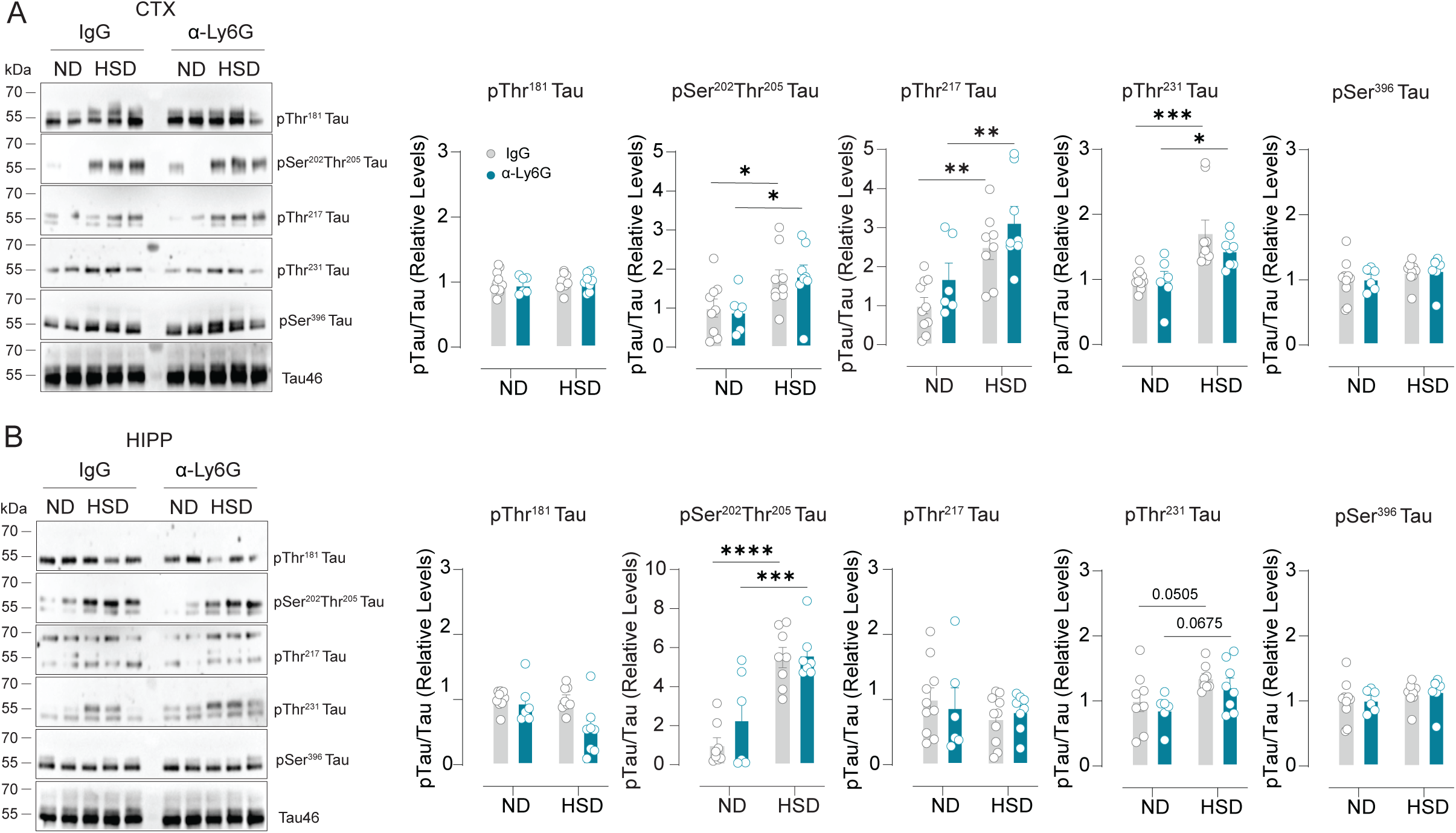
**Neutrophil depletion does not suppress the increase in phosphorylated tau induced by HSD**. **A.** HSD increases phosphorylation of tau on Ser^202^Thr^205^, Thr^217^ and Thr^231^ in the neocortex of both IgG control and anti-Ly6G-injected mice (Ser^202^Thr^205^, Diet: p=0.0030, Treatment: p=0.9889, Interaction: p=0.3628; Thr^217^, Diet: p=0.0001, Treatment: p=0.0544, Interaction: p=0.9263; Thr^231^, Diet: p=0.0001, Treatment: p=0.2868, Interaction: p=0.3725, n=6-10/group, two-way ANOVA plus Tukey’s multiple comparisons test). **B.** HSD increases phosphorylation of tau on Ser^202^Thr^205^ and Thr^231^ in the hippocampus of both IgG control and anti-Ly6G-injected mice (Ser^202^Thr^205^, Diet: p<0.0001, Treatment: p=0.2384, Interaction: p=0.3361; Thr^231^, Diet: p=0.0097, Treatment: p=0.3010, Interaction: p=0.9946, n=6-8/group, two-way ANOVA plus Tukey’s multiple comparisons test). Data is presented as mean ± SEM.

## DISCUSSION

This study provides the first evidence that brain capillary stalling is increased in mice fed a HSD. By using in vivo 2PEF microscopy, we found that capillary stalls were mostly caused by leukocytes and were in capillary branches proximal to the AV. Since the branches that were stalled had smaller vessel diameter, our data raise the possibility that vasoconstriction may contribute to the CBF reduction induced by HSD. Indeed, we found that RBCs velocity was reduced in upstream vessels including the PA. Administration of anti-Ly6G antibodies, depleted circulating neutrophils, reduced the number of stalled capillaries, and restored CBF in HSD mice, indicating that stalls may contribute to the reduction in CBF observed in these mice. In contrast, depletion of circulating neutrophils did not reduce brain p-Tau and did not improve the cognitive deficits associated with HSD. Altogether, these observations demonstrate that HSD promotes capillary stalling which, in turn, contributes to the CBF reduction observed in HSD mice. However, the CBF reduction mediated by capillary stalls does not play a role in the p-Tau accumulation and the cognitive impairment associated with HSD.

Our data are in contrast with previous observations indicating that restoring normal capillary flow improves cognitive function in mice models of Alzheimer’s disease (AD)^34^ and type-1 diabetes^43^. On the other hand, our findings are consistent with recent data demonstrating that reduction of capillary stalling increased CBF but did not improve memory deficits in wild-type mice fed a high-fat diet (HFD)^36^, indicating that hypoperfusion may not contribute to the cognitive impairment associated with HFD^66^. Furthermore, our findings are in line with previous data demonstrating that HSD-induced cognitive impairment is rescued by administration of anti-tau antibodies despite persistent reduction in CBF^44^, further indicating that CBF reduction is not sufficient for the cognitive impairment induced by HSD. Therefore, inasmuch as cerebrovascular alterations can mediate cognitive impairment^67^, these data suggest that factors other than hypoperfusion mediate the cognitive decline associated with cardiovascular risk factors^44^.

A key question concern how HSD promotes stalling of neutrophils in brain capillaries leading to CBF reduction. A critical step involves the interaction between leukocytes and ECs. Rolling and adhesion of neutrophils to the endothelium is a complex process that requires the expression of adhesion molecules on both neutrophils and ECs^68^. Rolling occurs even under resting conditions and is mainly mediated by selectin-mediated transient interactions^68^. However, in the presence of endothelial activation, increased expression of E-selectin enhances the interaction between ECs and neutrophils leading to a more stable adhesion through neutrophil LFA1 (CD11a/CD18) and MAC1 (CD11b/CD18) and endothelial ICAM1^68^. This scenario seems unlikely since our previous data indicate that HSD does not lead to expression of these inflammatory mediators on brain ECs^13^. Furthermore, in the present study we found that expression of adhesion molecules (i.e. *E-selectin, P-selectin, Icam1*) and neutrophil-chemoattractant molecules (i.e. *Cxcl1, Cxcl2*) were not increased in cerebral arteries or brain ECs isolated from HSD mice^13^, suggesting that brain EC activation may not play a role in the capillary stalling observed in HSD mice. Since HSD may alter the endothelial glycocalyx^69^, a possibility is that a thinned glycocalyx may increase neutrophil adhesion and capillary stalling^70^. In physiological conditions, the endothelial glycocalyx is negatively charged promoting the electrostatic repulsion of neutrophils^71^, and a damaged glycocalyx could promote adhesion of neutrophils to the endothelium^70^. Interestingly, pericyte loss has also been associated with alteration of the glycocalyx and capillary stalling^72^. However, since pericytes were not reduced in HSD mice^44^, this mechanism is unlikely to play a role. Another possibility is that reduced endothelial NO availability induced by HSD would promote leukocyte adhesion^30^. Specifically, NO might interfere with the ability of LFA1 and MAC1 to interact with the EC surface or it might suppress LFA1 and MAC1 expression on leukocytes as they roll along venular endothelium^30^. Nevertheless, this latter possibility seems unlikely since we found that, despite reduced endothelial NO availability^8–13^, markers of neutrophil activation, including CD18 and L-selectin, were not increased in brain neutrophils of HSD mice. These data are consistent with previous evidence demonstrating that exposure to high salt concentration does not activate neutrophils^73–76^. Thus, alternative mechanisms may mediate the interaction between neutrophils and brain ECs including the intriguing possibility that HSD may alter neutrophil glycoRNAs which have been recently emerged as crucial in the interaction between neutrophils and ECs^77^.

Finally, since stalling only affects a minority of brain capillaries in HSD mice, it is worth considering that, by analyzing all brain ECs and neutrophils, we might be missing subtle or localized effects that could only be observed by specifically looking at subsets of these cells. Indeed, the localization of the stalls indicate that ECs and/or neutrophil activation are confined to specific sections of the cerebrovascular tree. Thus, future studies investigating the effects of HSD at single cell-level could offer insights into the molecular and cellular mechanisms leading to stalling formation in HSD.

Systemic factors could also play a role in capillary stalling. Since HSD has been associated with increased secretion of von Willebrand factor by endothelial cells^78^, a possibility is that hypercoagulability and formation of intravascular clots may result in increased capillary stalling. Changes in the numbers of platelets and RBC could also play a role^33^. However, the finding that platelets are almost absent in stalled segments and that hematocrit was reduced in HSD mice (data not shown) exclude this possibility. Increased neutrophil counts may also contribute to stalling. In support of this hypothesis, the proinflammatory cytokine IL-17A, whose circulating levels are increased in HSD^13, 44^, has been shown to increase neutrophil counts via induction of G-CSF^79, 80^. However, the small increase in circulating neutrophils observed in HSD mice is unlikely to be crucial for stalls formation.

Another important finding of this study is that capillary stalling is associated with vasoconstriction of the stalled segment and CBF reduction. HSD reduces resting CBF and endothelium dependent relaxation^13, 25^ and the resulting vasoconstriction and slowing of capillary blood transit time could promote the formation of leukocyte and/or RBC capillary plugs. On the other hand, it could also be that the stalling promotes vasoconstriction. For example, stalled neutrophils may release reactive oxygen species (ROS) that could further scavenge NO and mediate vasoconstriction^81–86^. Although capillaries do not have SMC, ROS can induce pericyte contraction and capillary constriction^85, 86^. Furthermore, in an AD mouse model, ROS inhibition reduces capillary stalling, increases CBF and improves short-term memory^41^ indicating oxidative stress as critical for capillary stalling. Since neutrophil depletion rescues the vasoconstriction of the 1^st^ branch from the AV, the data support the possibility that neutrophil stalling may promote vasoconstriction. Future studies will have to examine further the complex interactions of microvascular flow, ROS, and capillary diameter in the development of capillary stalling.

In summary, our data demonstrated that HSD promotes stalling of neutrophils in brain capillaries and that depletion of circulating neutrophils can rescue the CBF reduction associated with HSD. However, depletion of circulating neutrophils did not reduce brain p-Tau and did not improve the cognitive deficits associated with HSD. Therefore, the CBF reduction mediated by stalling of neutrophils in brain capillaries is unlikely to be a key factor driving p-tau accumulation and cognitive impairment in HSD.

## FUNDING

This work was supported by the following grants: R01NS130045 (NINDS, GF), R01 R01NS095441 (NINDS, CI), CAF211776-01 (Cure Alzheimer’s Fund, GF and CI), BF A2023014F (BrightFocus Foundation, SA).

## AUTHOR CONTRIBUTIONS

SA, CI, and GF were responsible for the study design and for drafting and revising the manuscript. SA was responsible for the 2PFE studies and related analyses. BG was responsible for the behavioral studies. RS and LGB were responsible for the flow cytometry studies. BG, CB, and GR were responsible for the biochemical and molecular studies. LGB and JA provided critically input in the final version of the manuscript.

## DECLARATION OF CONFLICTING INTERESTS

C. Iadecola serves on the scientific advisory board of Broadview Ventures.

## DATA AVAILABILITY STATEMENT

The data that support the findings of this study are available from the corresponding author upon reasonable request.

## REFERENCES

1. Wang K, Jin Y, Wang M, Liu J, Bu X, Mu J, Lu J. Global cardiovascular diseases burden attributable to high sodium intake from 1990 to 2019. J Clin Hypertens (Greenwich) 2023;25:868–879.

2. Cook NR, Appel LJ, Whelton PK. Lower levels of sodium intake and reduced cardiovascular risk. Circulation 2014;129:981–989.

3. Tuomilehto J, Jousilahti P, Rastenyte D, Moltchanov V, Tanskanen A, Pietinen P, Nissinen A. Urinary sodium excretion and cardiovascular mortality in Finland: a prospective study. Lancet 2001;357:848–851.

4. Strazzullo P, D’Elia L, Kandala N-B, Cappuccio FP. Salt intake, stroke, and cardiovascular disease: meta-analysis of prospective studies. Bmj 2009;339:b4567.

5. Gardener H, Rundek T, Wright CB, Elkind MSV, Sacco RL. Dietary Sodium and Risk of Stroke in the Northern Manhattan Study. Stroke 2012;43:1200–1205.

6. Robinson AT, Edwards DG, Farquhar WB. The Influence of Dietary Salt Beyond Blood Pressure. Curr Hypertens Rep 2019;21:1–11.

7. Farquhar WB, Edwards DG, Jurkovitz CT, Weintraub WS. Dietary Sodium and Health: More Than Just Blood Pressure. J Am Coll Cardiol 2015;65:1042–1050.

8. Liu Y, Rusch NJ, Lombard JH. Loss of Endothelium and Receptor-Mediated Dilation in Pial Arterioles of Rats Fed a Short-Term High Salt Diet. Hypertension 1999;33:686–688.

9. Sylvester FA, Steep DW, Frisbee JC, Lombard JH. High-salt diet depresses acetylcholine reactivity proximal to NOS activation in cerebral arteries. Am J Physiol Heart Circ Physiol 2002;283:H353–H363.

10. Lombard JH, Sylvester FA, Phillips SA, Frisbee JC. High-salt diet impairs vascular relaxation mechanisms in rat middle cerebral arteries. Am J Physiol Heart Circ Physiol 2002;284:H1124–H1133.

11. Foulquier S, Dupuis F, Perrin-Sarrado C, Maguin Gate K, Merhi-Soussi F, Liminana P, Kwan YW, Capdeville-Atkinson C, Lartaud I, Atkinson J. High salt intake abolishes AT(2)-mediated vasodilation of pial arterioles in rats. J Hypertens 2011;29:1392–1399.

12. Durand MJ, Lombard JH. Low-dose angiotensin II infusion restores vascular function in cerebral arteries of high salt-fed rats by increasing copper/zinc superoxide dimutase expression. Am J Hypertens 2013;26:739–747.

13. Faraco G, Brea D, Garcia-Bonilla L, Wang G, Racchumi G, Chang H, Buendia I, Santisteban MM, Segarra SG, Koizumi K, Sugiyama Y, Murphy M, Voss H, Anrather J, Iadecola C. Dietary salt promotes neurovascular and cognitive dysfunction through a gut-initiated TH17 response. Nat Neurosci 2018;21:240–249.

14. Perry IJ, Beevers DG. Salt intake and stroke: a possible direct effect. J Hum Hypertens 1992;6:23–25.

15. Nagata C, Takatsuka N, Shimizu N, Shimizu H. Sodium intake and risk of death from stroke in Japanese men and women. Stroke 2004;35:1543–1547.

16. Strazzullo P, D’Elia L, Kandala NB, Cappuccio FP. Salt intake, stroke, and cardiovascular disease: meta-analysis of prospective studies. BMJ 2009;339:b4567.

17. Aaron KJ, Sanders PW. Role of dietary salt and potassium intake in cardiovascular health and disease: a review of the evidence. Mayo Clin Proc 2013;88:987–995.

18. O’Donnell M, Mente A, Rangarajan S, McQueen MJ, Wang X, Liu L, Yan H, Lee SF, Mony P, Devanath A, Rosengren A, Lopez-Jaramillo P, Diaz R, Avezum A, Lanas F, Yusoff K, Iqbal R, Ilow R, Mohammadifard N, Gulec S, Yusufali AH, Kruger L, Yusuf R, Chifamba J, Kabali C, Dagenais G, Lear SA, Teo K, Yusuf S. Urinary Sodium and Potassium Excretion, Mortality, and Cardiovascular Events. N Engl J Med 2014;371:612–623.

19. Okayama A, Okuda N, Miura K, Okamura T, Hayakawa T, Akasaka H, Ohnishi H, Saitoh S, Arai Y, Kiyohara Y, Takashima N, Yoshita K, Fujiyoshi A, Zaid M, Ohkubo T, Ueshima H, Group NDR. Dietary sodium-to-potassium ratio as a risk factor for stroke, cardiovascular disease and all-cause mortality in Japan: the NIPPON DATA80 cohort study. BMJ Open 2016;6:e011632.

20. Li Y, Huang Z, Jin C, Xing A, Liu Y, Huangfu C, Lichtenstein AH, Tucker KL, Wu S, Gao X. Longitudinal Change of Perceived Salt Intake and Stroke Risk in a Chinese Population. Stroke 2018;49:1332–1339.

21. Zhang J, Guo X, Lu Z, Tang J, Li Y, Xu A, Liu S. Cardiovascular Diseases Deaths Attributable to High Sodium Intake in Shandong Province, China. J Am Heart Assoc 2019;8:e010737.

22. He FJ, Tan M, Ma Y, MacGregor GA. Salt Reduction to Prevent Hypertension and Cardiovascular Disease: JACC State-of-the-Art Review. J Am Coll Cardiol 2020;75:632–647.

23. Makin SDJ, Mubki GF, Doubal FN, Shuler K, Staals J, Dennis MS, Wardlaw JM. Small Vessel Disease and Dietary Salt Intake: Cross-Sectional Study and Systematic Review. J Stroke Cerebrovasc Dis 2017;26:3020–3028.

24. Liu D, Zhang Q, Xing S, Wei F, Li K, Zhao Y, Zhang H, Gong G, Guo Y, Liu Z. Excessive salt intake accelerates the progression of cerebral small vessel disease in older adults. BMC Geriatr 2023;23:263.

25. Faraco G. Dietary salt, vascular dysfunction, and cognitive impairment. Cardiovasc Res 2024.

26. Mohan D, Yap KH, Reidpath D, Soh YC, McGrattan A, Stephan BCM, Robinson L, Chaiyakunapruk N, Siervo M, De PECt. Link Between Dietary Sodium Intake, Cognitive Function, and Dementia Risk in Middle-Aged and Older Adults: A Systematic Review. J Alzheimers Dis 2020;76:1347–1373.

27. Sindi S, Kåreholt I, Eskelinen M, Hooshmand B, Lehtisalo J, Soininen H, Ngandu T, Kivipelto M. Healthy Dietary Changes in Midlife Are Associated with Reduced Dementia Risk Later in Life. Nutrients 2018;10:1649–1611.

28. Blumenthal JA, Smith PJ, Mabe S, Hinderliter A, Lin PH, Liao L, Welsh-Bohmer KA, Browndyke JN, Kraus WE, Doraiswamy PM, Burke JR, Sherwood A. Lifestyle and neurocognition in older adults with cognitive impairments: A randomized trial. Neurology 2019;92:e212–e223.

29. Toda N, Ayajiki K, Okamura T. Cerebral blood flow regulation by nitric oxide: recent advances. Pharmacol Rev 2009;61:62–97.

30. Kubes P, Suzuki M, Granger DN. Nitric oxide: an endogenous modulator of leukocyte adhesion. Proc Natl Acad Sci U S A 1991;88:4651–4655.

31. Erdener SE, Dalkara T. Small Vessels Are a Big Problem in Neurodegeneration and Neuroprotection. Front Neurol 2019;10:889.

32. Ostergaard L. Blood flow, capillary transit times, and tissue oxygenation. The centennial of capillary recruitment. J Appl Physiol (1985) 2020;in press.

33. Santisakultarm TP, Paduano CQ, Stokol T, Southard TL, Nishimura N, Skoda RC, Olbricht WL, Schafer AI, Silver RT, Schaffer CB. Stalled cerebral capillary blood flow in mouse models of essential thrombocythemia and polycythemia vera revealed by in vivo two-photon imaging. J Thromb Haemost 2014;12:2120–2130.

34. Cruz Hernandez JC, Bracko O, Kersbergen CJ, Muse V, Haft-Javaherian M, Berg M, Park L, Vinarcsik LK, Ivasyk I, Rivera DA, Kang Y, Cortes-Canteli M, Peyrounette M, Doyeux V, Smith A, Zhou J, Otte G, Beverly JD, Davenport E, Davit Y, Lin CP, Strickland S, Iadecola C, Lorthois S, Nishimura N, Schaffer CB. Neutrophil adhesion in brain capillaries reduces cortical blood flow and impairs memory function in Alzheimer’s disease mouse models. Nat Neurosci 2019;22:413–420.

35. El Amki M, Gluck C, Binder N, Middleham W, Wyss MT, Weiss T, Meister H, Luft A, Weller M, Weber B, Wegener S. Neutrophils Obstructing Brain Capillaries Are a Major Cause of No-Reflow in Ischemic Stroke. Cell Rep 2020;33:108260.

36. Bracko O, Vinarcsik LK, Cruz Hernandez JC, Ruiz-Uribe NE, Haft-Javaherian M, Falkenhain K, Ramanauskaite EM, Ali M, Mohapatra A, Swallow MA, Njiru BN, Muse V, Michelucci PE, Nishimura N, Schaffer CB. High fat diet worsens Alzheimer’s disease-related behavioral abnormalities and neuropathology in APP/PS1 mice, but not by synergistically decreasing cerebral blood flow. Sci Rep 2020;10:9884.

37. Falkenhain K, Ruiz-Uribe NE, Haft-Javaherian M, Ali M, Stall C, Michelucci PE, Schaffer CB, Bracko O. A pilot study investigating the effects of voluntary exercise on capillary stalling and cerebral blood flow in the APP/PS1 mouse model of Alzheimer’s disease. PLoS One 2020;15:e0235691.

38. Rolfes L, Riek-Burchardt M, Pawlitzki M, Minnerup J, Bock S, Schmidt M, Meuth SG, Gunzer M, Neumann J. Neutrophil granulocytes promote flow stagnation due to dynamic capillary stalls following experimental stroke. Brain Behav Immun 2021;93:322–330.

39. Erdener SE, Tang J, Kilic K, Postnov D, Giblin JT, Kura S, Chen IA, Vayisoglu T, Sakadzic S, Schaffer CB, Boas DA. Dynamic capillary stalls in reperfused ischemic penumbra contribute to injury: A hyperacute role for neutrophils in persistent traffic jams. J Cereb Blood Flow Metab 2021;41:236–252.

40. Ali M, Falkenhain K, Njiru BN, Murtaza-Ali M, Ruiz-Uribe NE, Haft-Javaherian M, Catchers S, Nishimura N, Schaffer CB, Bracko O. VEGF signalling causes stalls in brain capillaries and reduces cerebral blood flow in Alzheimer’s mice. Brain 2022;145:1449–1463.

41. Ruiz-Uribe NE, Bracko O, Swallow M, Omurzakov A, Dash S, Uchida H, Xiang D, Haft-Javaherian M, Falkenhain K, Lamont ME, Ali M, Njiru BN, Chang HY, Tan AY, Xiang JZ, Iadecola C, Park L, Sanchez T, Nishimura N, Schaffer CB. Vascular oxidative stress causes neutrophil arrest in brain capillaries, leading to decreased cerebral blood flow and contributing to memory impairment in a mouse model of Alzheimeraeuros disease. bioRxiv 2023.

42. Lim HK, Bae S, Han K, Kang BM, Jeong Y, Kim SG, Suh M. Seizure-induced neutrophil adhesion in brain capillaries leads to a decrease in postictal cerebral blood flow. iScience 2023;26:106655.

43. Sharma S, Cheema M, Reeson PL, Narayana K, Boghozian R, Cota AP, Brosschot TP, FitzPatrick RD, Korbelin J, Reynolds LA, Brown CE. A pathogenic role for IL-10 signalling in capillary stalling and cognitive impairment in type 1 diabetes. Nat Metab 2024.

44. Faraco G, Hochrainer K, Segarra SG, Schaeffer S, Santisteban MM, Menon A, Jiang H, Holtzman DM, Anrather J, Iadecola C. Dietary salt promotes cognitive impairment through tau phosphorylation. Nature 2019;574:686–690.

45. DeVos SL, Hyman BT. Tau at the Crossroads between Neurotoxicity and Neuroprotection. Neuron 2017;94:703–704.

46. Raz L, Bhaskar K, Weaver J, Marini S, Zhang Q, Thompson JF, Espinoza C, Iqbal S, Maphis NM, Weston L, Sillerud LO, Caprihan A, Pesko JC, Erhardt EB, Rosenberg GA. Hypoxia promotes tau hyperphosphorylation with associated neuropathology in vascular dysfunction. Neurobiology of Disease 2018:1–0.

47. Zhao Y, Gu JH, Dai CL, Liu Q, Iqbal K, Liu F, Gong CX. Chronic cerebral hypoperfusion causes decrease of O-GlcNAcylation, hyperphosphorylation of tau and behavioral deficits in mice. Front Aging Neurosci 2014;6:10.

48. Antoniou X, Gassmann M, Ogunshola OO. Cdk5 interacts with Hif-1alpha in neurons: a new hypoxic signalling mechanism? Brain Res 2011;1381:1–10.

49. Schager B, Brown CE. Susceptibility to capillary plugging can predict brain region specific vessel loss with aging. J Cereb Blood Flow Metab 2020;40:2475–2490.

50. Zhao Y, Gao P, Sun F, Li Q, Chen J, Yu H, Li L, Wei X, He H, Lu Z, Wei X, Wang B, Cui Y, Xiong S, Shang Q, Xu A, Huang Y, Liu D, Zhu Z. Sodium Intake Regulates Glucose Homeostasis through the PPARdelta/Adiponectin-Mediated SGLT2 Pathway. Cell Metab 2016;23:699–711.

51. Powles J, Fahimi S, Micha R, Khatibzadeh S, Shi P, Ezzati M, Engell RE, Lim SS, Danaei G, Mozaffarian D, Global Burden of Diseases N, Chronic Diseases Expert G. Global, regional and national sodium intakes in 1990 and 2010: a systematic analysis of 24 h urinary sodium excretion and dietary surveys worldwide. BMJ Open 2013;3:e003733.

52. Kang L, Yu H, Yang X, Zhu Y, Bai X, Wang R, Cao Y, Xu H, Luo H, Lu L, Shi MJ, Tian Y, Fan W, Zhao BQ. Neutrophil extracellular traps released by neutrophils impair revascularization and vascular remodeling after stroke. Nat Commun 2020;11:2488.

53. Santisteban MM, Ahn SJ, Lane D, Faraco G, Garcia-Bonilla L, Racchumi G, Poon C, Schaeffer S, Segarra SG, Korbelin J, Anrather J, Iadecola C. Endothelium-Macrophage Crosstalk Mediates Blood-Brain Barrier Dysfunction in Hypertension. Hypertension 2020;76:795–807.

54. Schaffer CB, Friedman B, Nishimura N, Schroeder LF, Tsai PS, Ebner FF, Lyden PD, Kleinfeld D. Two-photon imaging of cortical surface microvessels reveals a robust redistribution in blood flow after vascular occlusion. PLoS Biol 2006;4:e22.

55. Santisakultarm TP, Cornelius NR, Nishimura N, Schafer AI, Silver RT, Doerschuk PC, Olbricht WL, Schaffer CB. In vivo two-photon excited fluorescence microscopy reveals cardiac-and respiration-dependent pulsatile blood flow in cortical blood vessels in mice. Am J Physiol Heart Circ Physiol 2012;302:H1367–1377.

56. Benakis C, Brea D, Caballero S, Faraco G, Moore J, Murphy M, Sita G, Racchumi G, Ling L, Pamer EG, Iadecola C, Anrather J. Commensal microbiota affects ischemic stroke outcome by regulating intestinal gammadelta T cells. Nat Med 2016;22:516–523.

57. Livak KJ, Schmittgen TD. Analysis of relative gene expression data using real-time quantitative PCR and the 2(-Delta Delta C(T)) Method. Methods 2001;25:402–408.

58. Deacon RM. Assessing nest building in mice. Nat Protoc 2006;1:1117–1119.

59. Faraco G, Sugiyama Y, Lane D, Garcia-Bonilla L, Chang H, Santisteban MM, Racchumi G, Murphy M, Van Rooijen N, Anrather J, Iadecola C. Perivascular macrophages mediate the neurovascular and cognitive dysfunction associated with hypertension. J Clin Invest 2016;126:4674–4689.

60. Cruz-Hernández JC, Bracko O, Kersbergen CJ, Muse V, Haft-Javaherian M, Berg M, Park L, Vinarcsik LK, Ivasyk I, Rivera DA, Kang Y, Cortes-Canteli M, Peyrounette M, Doyeux V, Smith A, Zhou J, Otte G, Beverly JD, Davenport E, Davit Y, Lin CP, Strickland S, Iadecola C, Lorthois S, Nishimura N, Schaffer CB. Neutrophil adhesion in brain capillaries reduces cortical blood flow and impairs memory function in Alzheimer’s disease mouse models. Nat Neurosci 2019;22:413–420.

61. Santizo RA, Xu HL, Galea E, Muyskens S, Baughman VL, Pelligrino DA. Combined endothelial nitric oxide synthase upregulation and caveolin-1 downregulation decrease leukocyte adhesion in pial venules of ovariectomized female rats. Stroke 2002;33:613–616.

62. Erdener SE, Tang J, Sajjadi A, Kilic K, Kura S, Schaffer CB, Boas DA. Spatio-temporal dynamics of cerebral capillary segments with stalling red blood cells. J Cereb Blood Flow Metab 2019;39:886–900.

63. Rahman I, Sanchez AC, Davies J, Rzeniewicz K, Abukscem S, Joachim J, Hoskins Green HL, Killock D, Sanz MJ, Charras G, Parsons M, Ivetic A. Correction: L-selectin regulates human neutrophil transendothelial migration. J Cell Sci 2022;135.

64. Stadtmann A, Germena G, Block H, Boras M, Rossaint J, Sundd P, Lefort C, Fisher CI, Buscher K, Gelschefarth B, Urzainqui A, Gerke V, Ley K, Zarbock A. The PSGL-1-L-selectin signaling complex regulates neutrophil adhesion under flow. J Exp Med 2013;210:2171–2180.

65. Garcia-Bonilla L, Shahanoor Z, Sciortino R, Nazarzoda O, Racchumi G, Iadecola C, Anrather J. Analysis of brain and blood single-cell transcriptomics in acute and subacute phases after experimental stroke. Nat Immunol 2024;25:357–370.

66. Zuloaga KL, Johnson LA, Roese NE, Marzulla T, Zhang W, Nie X, Alkayed FN, Hong C, Grafe MR, Pike MM, Raber J, Alkayed NJ. High fat diet-induced diabetes in mice exacerbates cognitive deficit due to chronic hypoperfusion. J Cereb Blood Flow Metab 2016;36:1257–1270.

67. Iadecola C, Duering M, Hachinski V, Joutel A, Pendlebury ST, Schneider JA, Dichgans M. Vascular Cognitive Impairment and Dementia: JACC Scientific Expert Panel. J Am Coll Cardiol 2019;73:3326–3344.

68. Nemeth T, Sperandio M, Mocsai A. Neutrophils as emerging therapeutic targets. Nat Rev Drug Discov 2020;19:253–275.

69. Oberleithner H, Peters W, Kusche-Vihrog K, Korte S, Schillers H, Kliche K, Oberleithner K. Salt overload damages the glycocalyx sodium barrier of vascular endothelium. Pflugers Arch 2011;462:519–528.

70. Yoon JH, Shin P, Joo J, Kim GS, Oh WY, Jeong Y. Increased capillary stalling is associated with endothelial glycocalyx loss in subcortical vascular dementia. J Cereb Blood Flow Metab 2022;42:1383–1397.

71. Reitsma S, Slaaf DW, Vink H, van Zandvoort MA, oude Egbrink MG. The endothelial glycocalyx: composition, functions, and visualization. Pflugers Arch 2007;454:345–359.

72. Choe YG, Yoon JH, Joo J, Kim B, Hong SP, Koh GY, Lee DS, Oh WY, Jeong Y. Pericyte Loss Leads to Capillary Stalling Through Increased Leukocyte-Endothelial Cell Interaction in the Brain. Front Cell Neurosci 2022;16:848764.

73. Nadesalingam A, Chen JHK, Farahvash A, Khan MA. Hypertonic Saline Suppresses NADPH Oxidase-Dependent Neutrophil Extracellular Trap Formation and Promotes Apoptosis. Front Immunol 2018;9:359.

74. Jobin K, Stumpf NE, Schwab S, Eichler M, Neubert P, Rauh M, Adamowski M, Babyak O, Hinze D, Sivalingam S, Weisheit C, Hochheiser K, Schmidt SV, Meissner M, Garbi N, Abdullah Z, Wenzel U, Holzel M, Jantsch J, Kurts C. A high-salt diet compromises antibacterial neutrophil responses through hormonal perturbation. Sci Transl Med 2020;12.

75. Krampert L, Bauer K, Ebner S, Neubert P, Ossner T, Weigert A, Schatz V, Toelge M, Schroder A, Herrmann M, Schnare M, Dorhoi A, Jantsch J. High Na(+) Environments Impair Phagocyte Oxidase-Dependent Antibacterial Activity of Neutrophils. Front Immunol 2021;12:712948.

76. Zlatar L, Mahajan A, Munoz-Becerra M, Weidner D, Bila G, Bilyy R, Titze J, Hoffmann MH, Schett G, Herrmann M, Steffen U, Munoz LE, Knopf J. Suppression of neutrophils by sodium exacerbates oxidative stress and arthritis. Front Immunol 2023;14:1174537.

77. Zhang N, Tang W, Torres L, Wang X, Ajaj Y, Zhu L, Luan Y, Zhou H, Wang Y, Zhang D, Kurbatov V, Khan SA, Kumar P, Hidalgo A, Wu D, Lu J. Cell surface RNAs control neutrophil recruitment. Cell 2024;187:846–860 e817.

78. Dmitrieva NI, Burg MB. Secretion of von Willebrand factor by endothelial cells links sodium to hypercoagulability and thrombosis. Proc Natl Acad Sci U S A 2014;111:6485–6490.

79. Schwarzenberger P, La Russa V, Miller A, Ye P, Huang W, Zieske A, Nelson S, Bagby GJ, Stoltz D, Mynatt RL, Spriggs M, Kolls JK. IL-17 stimulates granulopoiesis in mice: use of an alternate, novel gene therapy-derived method for in vivo evaluation of cytokines. J Immunol 1998;161:6383–6389.

80. Forlow SB, Schurr JR, Kolls JK, Bagby GJ, Schwarzenberger PO, Ley K. Increased granulopoiesis through interleukin-17 and granulocyte colony-stimulating factor in leukocyte adhesion molecule-deficient mice. Blood 2001;98:3309–3314.

81. Didion SP, Faraci FM. Angiotensin II produces superoxide-mediated impairment of endothelial function in cerebral arterioles. Stroke 2003;34:2038–2042.

82. Girouard H, Park L, Anrather J, Zhou P, Iadecola C. Angiotensin II attenuates endothelium-dependent responses in the cerebral microcirculation through nox-2-derived radicals. Arterioscler Thromb Vasc Biol 2006;26:826–832.

83. Faraci FM, Lamping KG, Modrick ML, Ryan MJ, Sigmund CD, Didion SP. Cerebral vascular effects of angiotensin II: new insights from genetic models. J Cereb Blood Flow Metab 2006;26:449–455.

84. Chrissobolis S, Faraci FM. Sex differences in protection against angiotensin II-induced endothelial dysfunction by manganese superoxide dismutase in the cerebral circulation. Hypertension 2010;55:905–910.

85. Nortley R, Korte N, Izquierdo P, Hirunpattarasilp C, Mishra A, Jaunmuktane Z, Kyrargyri V, Pfeiffer T, Khennouf L, Madry C, Gong H, Richard-Loendt A, Huang W, Saito T, Saido TC, Brandner S, Sethi H, Attwell D. Amyloid beta oligomers constrict human capillaries in Alzheimer’s disease via signaling to pericytes. Science 2019;365.

86. Korte N, Barkaway A, Wells J, Freitas F, Sethi H, Andrews SP, Skidmore J, Stevens B, Attwell D. Inhibiting Ca(2+) channels in Alzheimer’s disease model mice relaxes pericytes, improves cerebral blood flow and reduces immune cell stalling and hypoxia. Nat Neurosci 2024;27:2086–2100.

