## Supplementary figures and images for "Neutrophil stalling does not mediate the increase in tau phosphorylation and the cognitive impairment associated with high salt diet"

### Suppl. Figure 1

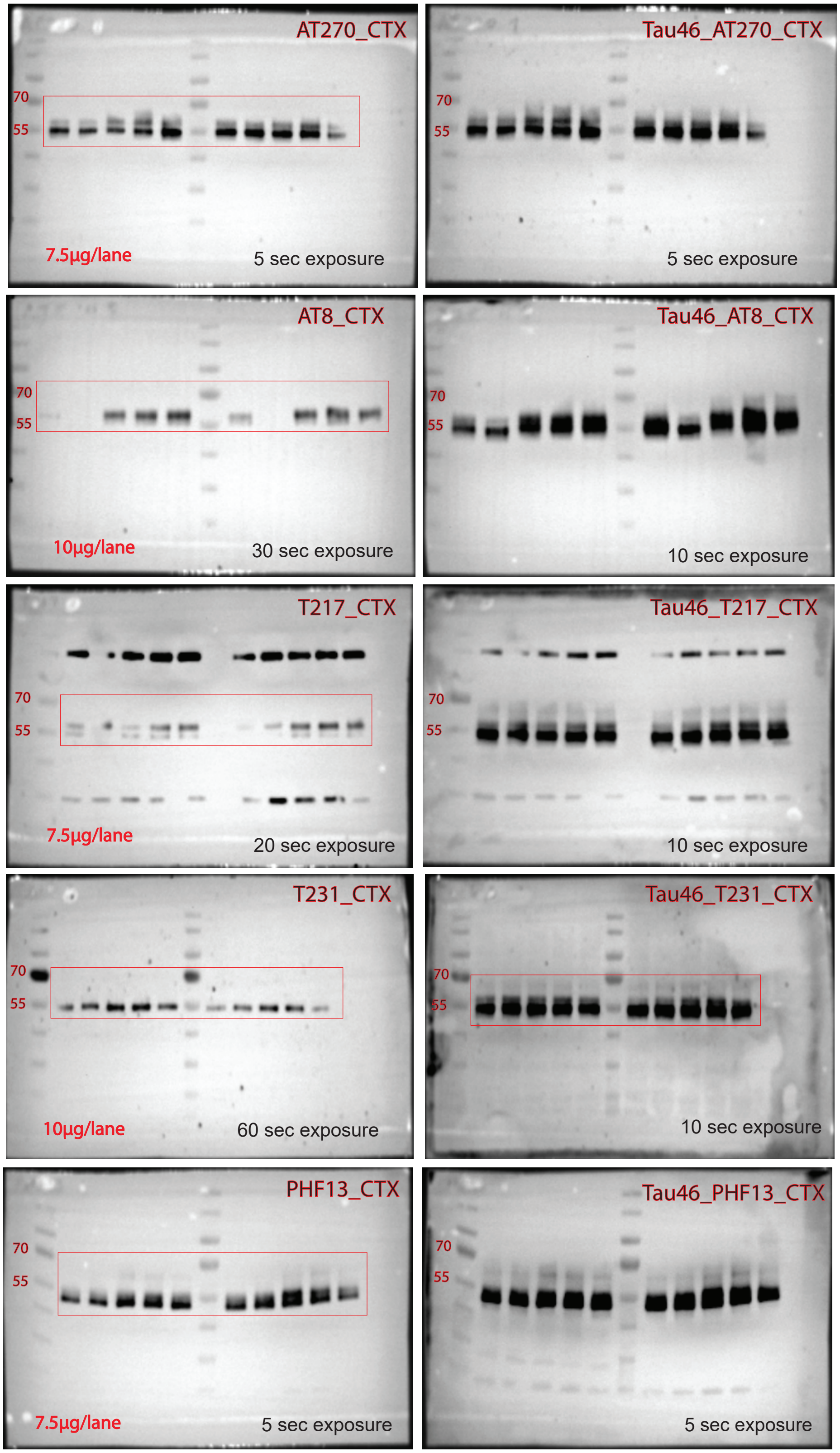


PageRuler™ Prestained Protein Ladder, 10 to 180 kDa, ThermoFisher #26616

### Suppl. Figure 2

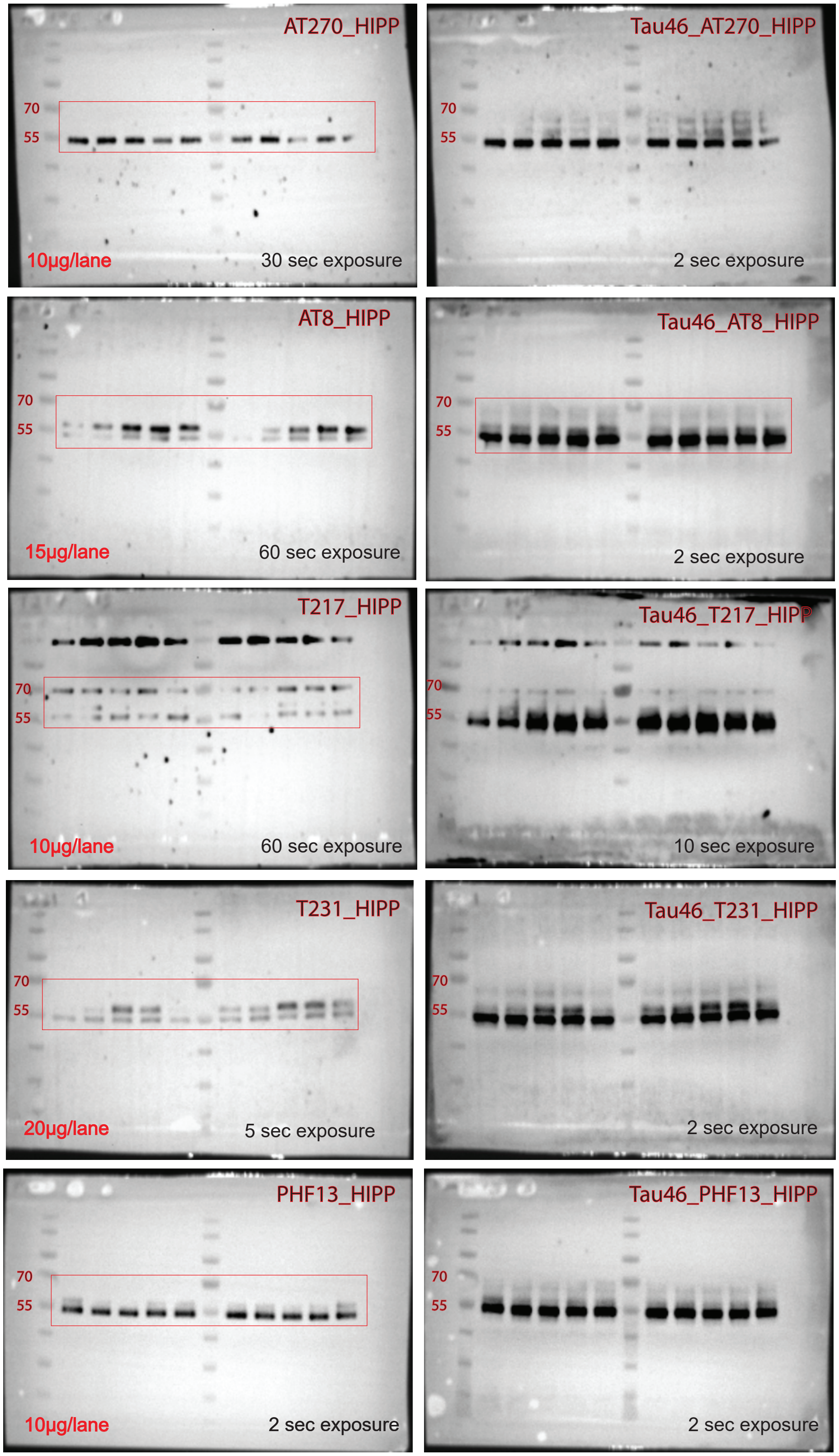


PageRuler™ Prestained Protein Ladder, 10 to 180 kDa, ThermoFisher #26616
