## Supplementary material for "Neutrophil stalling does not mediate the increase in tau phosphorylation and the cognitive impairment associated with high salt diet": Table 1

| *Cybb (Gp91)* | AATCTCAGGCCAATCACTTTG | AACGCCTATTGTGGTGTTAGG |
| --- | --- | --- |
| *Il1b* | CTCTCCACCTCAATGGACAGA | TTTTGTCGTTGCTTGGTTCTC |
| *Mmp9* | ATTCGCGTGGATAAGGAGTTC | CCTTGTTCACCTCATTTTGGA |
| *Cxcl2* | TGAACAAAGGCAAGGCTAACTG | GAGGCACATCAGGTACGATCC |
| *Hprt* | AGTGTTGGATACAGGCCAGAC | CGTGATTCAAATCCCTGAAGT |

**Table 1. List of primers used for RT-PCR studies on circulating neutrophils.**
