## Supplementary material for "Neutrophil stalling does not mediate the increase in tau phosphorylation and the cognitive impairment associated with high salt diet": Table 2

| **Name** | **Description** | **Source** | **Catalog No.** | **Dilution** |
| --- | --- | --- | --- | --- |
| pTau Ser^202^Thr^205^ (AT8) | Mouse mAb | ThermoFisher | MN1020 | 1:1000 |
| pTau Thr^217^ | Rabbit pAb | ThermoFisher | 44-744 | 1:1000 |
| pTau Ser^231^ (AT180) | Mouse mAb | ThermoFisher | MN1040 | 1:500 |
| pThr^231^ (RZ3) | Mouse mAb | Peter Davies | n/a | 1:500 |
| pTau Ser^396^ | Mouse mAb | Cell Signaling | 9632 | 1:1000 |
| pTau Thr^181^ (AT270) | Rabbit mAb | Cell Signaling | 12885 | 1:1000 |
| Tau46 | Mouse mAb | Cell Signaling | 4019 | 1:1000 |

**Table 2. List of primary antibodies used for detection of phosphorylated tau.**
